# A Community Challenge to Predict Clinical Outcomes After Immune Checkpoint Blockade in Non-Small Cell Lung Cancer

**DOI:** 10.1101/2022.12.05.518667

**Authors:** Mike Mason, Óscar Lapuente-Santana, Anni S. Halkola, Wenyu Wang, Raghvendra Mall, Xu Xiao, Jacob Kaufman, Jingxin Fu, Jacob Pfeil, Jineta Banerjee, Verena Chung, Han Chang, Scott D. Chasalow, Hung Ying Lin, Rongrong Chai, Thomas Yu, Francesca Finotello, Tuomas Mirtti, Mikko I. Mäyränpää, Jie Bao, Emmy W. Verschuren, Eiman I. Ahmed, Michele Ceccarelli, Lance D. Miller, Gianni Monaco, Wouter R.L. Hendrickx, Shimaa Sherif, Lin Yang, Ming Tang, Shengqing Stan Gu, Wubing Zhang, Yi Zhang, Zexian Zeng, Avinash Das Sahu, Yang Liu, Wenxian Yang, Davide Bedognetti, Jing Tang, Federica Eduati, Teemu D. Laajala, William J. Geese, Justin Guinney, Joseph D. Szustakowski, David P. Carbone, Benjamin G. Vincent

**Author notes:** Lead authors from participating teams in the Anti–PD-1 Response Prediction DREAM Challenge with equal contribution. Senior authors from participating teams in the Anti–PD-1 Response Prediction DREAM Challenge with equal contribution. Co-senior authors from the Anti–PD-1 Response Prediction DREAM Challenge steering committee. **Corresponding author’s contact details:** Mike Mason. **Previous presentation:** Results from this study have been presented, in part, at the 14^th^ annual RECOMB/ISCB Conference on Regulatory and Systems Genomics with DREAM Challenges RSGDREAM 2022), November 8–9, 2022, Las Vegas, Nevada, USA.

## Abstract

**Purpose:** Predictive biomarkers of immune checkpoint inhibitors (ICIs) efficacy are currently lacking for non-small cell lung cancer (NSCLC). Here, we describe the results from the Anti–PD-1 Response Prediction DREAM Challenge, a crowdsourced initiative that enabled the assessment of predictive models by using data from two randomized controlled clinical trials (RCTs) of ICIs in first-line metastatic NSCLC.

**Methods:** Participants developed and trained models using public resources. These were evaluated with data from the CheckMate 026 trial (NCT02041533), according to the model-to-data paradigm to maintain patient confidentiality. The generalizability of the models with the best predictive performance was assessed using data from the CheckMate 227 trial (NCT02477826). Both trials were phase III RCTs with a chemotherapy control arm, which supported the differentiation between predictive and prognostic models. Isolated model containers were evaluated using a bespoke strategy that considered the challenges of handling transcriptome data from clinical trials.

**Results:** A total of 59 teams participated, with 417 models submitted. Multiple predictive models, as opposed to a prognostic model, were generated for predicting overall survival, progression-free survival, and progressive disease status with ICIs. Variables within the models submitted by participants included tumor mutational burden (TMB), programmed death ligand 1 (PD-L1) expression, and gene-expression–based signatures. The best-performing models showed improved predictive power over reference variables, including TMB or PD-L1.

**Conclusion:** This DREAM Challenge is the first successful attempt to use protected phase III clinical data for a crowdsourced effort towards generating predictive models for ICIs clinical outcomes and could serve as a blueprint for similar efforts in other tumor types and disease states, setting a benchmark for future studies aiming to identify biomarkers predictive of ICIs efficacy.

**Context summary:** *Key objective:* Not all patients with non-small cell lung cancer (NSCLC) eligible for immune checkpoint inhibitor (ICIs) respond to treatment, but accurate predictive biomarkers of ICIs clinical outcomes are currently lacking. This crowdsourced initiative enabled the robust assessment of predictive models using data from two randomized clinical trials of first-line ICI in metastatic NSCLC.

*Knowledge generated:* Models submitted indicate that a combination of programmed death ligand 1 (PD-L1), tumor mutational burden (TMB), and immune gene signatures might be able to identify patients more likely to respond to ICIs. TMB and PD-L1 seemed important to predict progression-free survival and overall survival. Mechanisms including apoptosis, T-cell crosstalk, and adaptive immune resistance appeared essential to predict response.

*Relevance:* 

## Introduction

Immune checkpoint inhibitors (ICIs) have revolutionized cancer treatment, with advanced non-small cell lung cancer (NSCLC) among the tumor types showing longer survival with ICIs than with chemotherapy in multiple treatment lines.^1–4^ While ICIs have demonstrated high response rates in some tumor types,^5^ not all patients with advanced cancer eligible for ICIs respond to them, highlighting the need for biomarkers predictive of their efficacy.^6–8^

Multiple biomarkers have been explored as predictors of clinical outcomes, including programmed death ligand 1 (PD-L1) expression and tumor mutational burden (TMB), which are used in clinical practice but are imperfect predictors of ICI response and not standardized across studies.^9^ Associations between clinical outcomes with ICIs and certain biomarkers, including immune-related gene expression, gene signatures, and adaptive immune receptor repertoire features (eg, T-cell–inflamed gene expression, chemokine expression, immunologic constant of rejection [ICR], T-cell receptor repertoire clonality) have been reported.^10–14^ However, a comparison of performance of these markers using large, independent validation datasets is lacking. Biomarker studies in NSCLC have been limited by small sample sizes and lack of a chemotherapy control arm, preventing differentiation between prognostic and predictive biomarkers.^15–18^ Robust predictive biomarkers will be critical to identify who would be more likely to benefit from ICIs, and could guide treatment choice and serve as trial stratification factors.

Here, we describe the Anti–PD-1 Response Prediction DREAM Challenge, a crowdsourced initiative that enabled the assessment of predictive models using data from two randomized clinical trials (RCTs) of first-line ICIs in NSCLC. We used an innovative model-to-data paradigm that enabled broad participation without requiring direct access to restricted data. This approach protected patient confidentiality while mitigating the risk of overfitting, lack of replicability, and irreproducibility.^19,20^

The pioneering design of this Challenge addressed scientific and technical issues that the community has faced in identifying robust predictors of ICI efficacy. The engagement of worldwide researchers using a reference dataset and consistent metrics leveled the playing field and allowed for head-to-head comparisons of model performance. The use of data from large, mature, well-annotated RCTs eliminated, at least partially, the limitations of analyses based on smaller trials, observational studies, or restricted sample cohorts. Metrics using information from both treatment and control arms allow the differentiation of prognostic models from those that are predictive of population-level benefit from ICI therapies. Finally, the combination of closed competitive and open cooperative phases of this Challenge enabled unprecedented collaboration among academic and industry leaders.

## Materials and Methods

### Challenge Questions

A steering committee, including members from Bristol Myers Squibb, Sage Bionetworks, and oncology physician-scientists, developed clinically relevant questions that could be addressed through the DREAM Challenge framework. This Challenge comprised three sub-challenges to identify models predictive of progression-free survival (PFS), overall survival (OS), and best overall response (BOR) of progressive disease (PD) with ICI treatment (Table 1).^21^

**TABLE 1.**
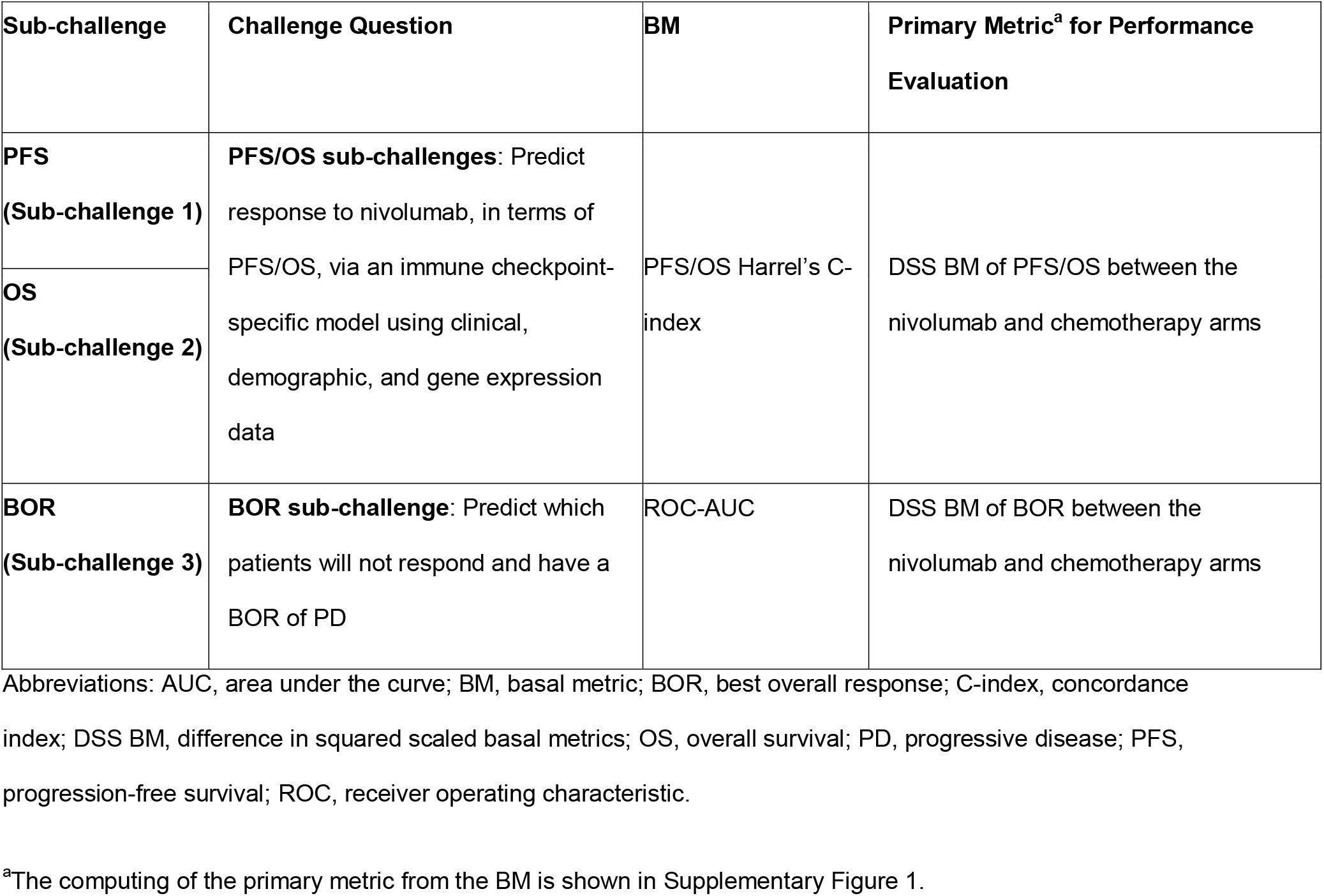
Challenge Questions and Metrics Used for Performance Evaluation^21^.

### Training and Validation Datasets

The design of the Challenge is summarized in Figure 1. To protect patient confidentiality, participants could not access directly the evaluation dataset (CheckMate 026), in line with the model-to-data paradigm.^19^ Because of the abundance of publicly available datasets, participants were not provided training data, thereby maintaining a large testing dataset. The variables available to participants and details on the training data used for model construction are shown in Supplementary Table 1 and Supplementary Methods 1, respectively. Gene-expression–based predictors are shown in Supplementary Tables 2 and 3. Participants developed and trained predictive models using publicly available resources, including those referenced on the Challenge website (TIDE resources,^22^ The Cancer Research Institute’s iAtlas,^23^ and other published data)^24^ and other datasets accessible via their institutions. To ensure proper execution of the independently trained models on the embargoed evaluation dataset, a synthetic dataset with the same formatting as the evaluation dataset was available. Participants submitted dockerized models^25^ consisting of the model itself plus software components to run the model in the DREAM evaluation infrastructure (Supplementary Methods 2). This approach supported reproducibility and a platform-independent evaluation of submitted models. Each team could submit different models for each sub-challenge.

**FIG 1.**
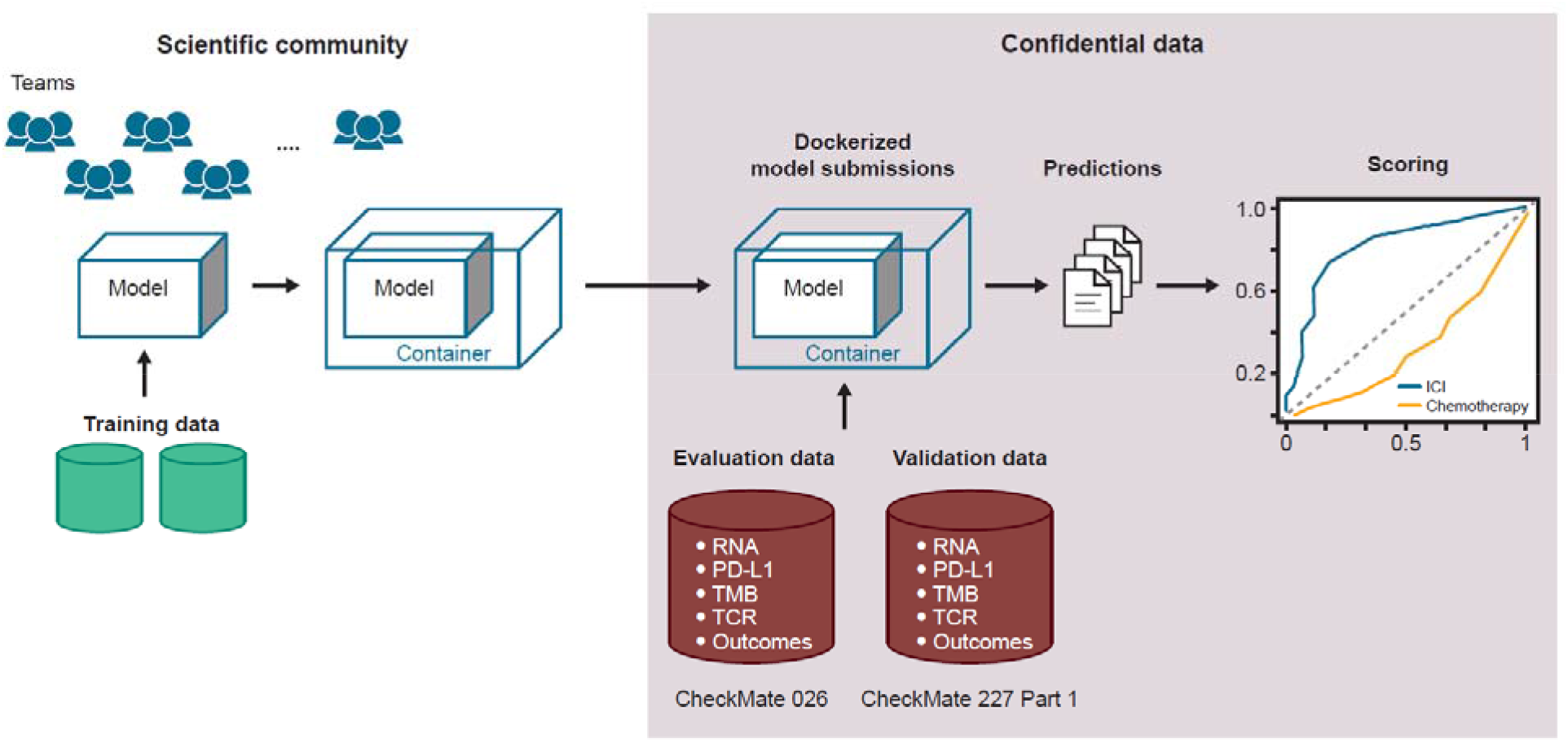
Challenge design. ICI, immune checkpoint inhibitor; PD-L1, programmed death ligand 1; TCR, T-cell receptor; TMB, tumor mutational burden.

The evaluation dataset from CheckMate 026 (NCT02041533)^26^ was selected because it was large, contained multimodal data, was well-characterized at the clinical and molecular level, and allowed potential differentiation between predictive and prognostic models.^27^ In CheckMate 026, patients with untreated stage IV or recurrent NSCLC and tumor PD-L1 ≥ 1% were randomized 1:1 to receive nivolumab or platinum-based chemotherapy.^26^ Top-performing models identified with CheckMate 026 data were validated on an independent dataset from CheckMate 227 (Part 1) (NCT02477826) in patients with stage IV or recurrent NSCLC.^28,29^ Identification of potential biomarkers of response to nivolumab were protocol-defined exploratory end points in both CheckMate 026 and 227. In CheckMate 227, patients with tumor PD-L1 ≥ 1% (Part 1a) received either nivolumab + ipilimumab, nivolumab monotherapy, or chemotherapy; patients with PD-L1 < 1% (Part 1b) received either nivolumab + ipilimumab, nivolumab + chemotherapy, or chemotherapy for the first-line treatment of metastatic NSCLC.^28,29^ Top-performing models were validated in the nivolumab + ipilimumab arms of CheckMate 227 in patients with any level of PD-L1 expression, as these arms were part of the successful primary end points of that trial. Baseline characteristics of patients in CheckMate 026 and 227 were published previously (Supplementary Tables 4 and 5).^26,28,29^

### Assessing Model Performance

Performance metrics (Table 1) were designed to identify predictive rather than prognostic models: top-performing models should accurately rank response measures for patients in the ICI arm but not in the chemotherapy arm to reflect a model’s capacity to inform a clinical decision in favor of one therapy over another. For the PFS sub-challenge, we computed Harrell’s concordance index (C-index) of PFS and model predictions as a basal metric (BM) calculated in each arm.^30^ We used the C-index in the OS sub-challenge after first correcting for potential effects caused by patient crossover from the chemotherapy arm to the nivolumab arm in CheckMate 026.^31^ For the BOR sub-challenge, the BM was the receiver operating characteristic (ROC) area under the curve (AUC) of the model predictions in each arm.

For each sub-challenge, the primary metric (DSS) was the difference in squared scaled BM between the nivolumab arm and chemotherapy arm, where *scaled* (*BM*) = 2 × (*BM* − 0.5) (Table 1, Supplementary Figure 1).^32,33^ Models that performed well in the nivolumab arm and randomly in the chemotherapy arm had positive primary scores. Models that performed well in the chemotherapy arm but randomly in the nivolumab arm had negative primary scores. Models that performed the same in each arm had a score of 0. Squaring of the BM allowed us to accommodate models that predicted well in the negative direction as good predictors.

A team’s model performance was determined in each sub-challenge. To be eligible for top-performing status, a model had to outperform the TMB baseline model based on the primary metric (Bayes factor relative to TMB baseline model, *K_TMB_* > 3, see Supplementary Methods 2). A description of baseline models and published reference models is provided in Supplementary Tables 2 and 3. For models meeting this criterion, we computed *K_DSS_Max_*, the Bayes factor relative to the highest primary metric in that sub-challenge. Models with *K_DSS_Max_* < 3 were considered tied with the highest scoring model. The BM from the nivolumab arm was used for tie-breaking. If multiple tied models had tie-breaking scores close to the best tie-breaking score, they were included as top-performers for the sub-challenge.

## Results

### Overall Participation in This Challenge

Fifty-one teams and eight individuals made at least one valid submission to the Challenge, with 417 models submitted across the three sub-challenges. Top-performing model descriptions are available on the Challenge website, Supplementary Methods 1, and Table 2. Author teams’ contributions to their respective model are reported in the author teams’ contribution section of the Supplement. Top-performing models outperformed the 14 comparator models for each sub-challenge.

**TABLE 2.** Description of Top-Performing Models.

| Model Name | Model Description |
| --- | --- |
| **Aginome-Amoy**<br>**Top-performer in the BOR sub-challenge** | A rule-based model was generated using patients stratified into three groups based on their PD-L1 and TMB expression scores:<br>Group 1: PD-L1 score below median<br>Group 2: PD-L1 score above median and TMB score below median<br>Group 3: Both PD-L1 and TMB expression scores above median<br>The following heuristic rules were used to decide the ranking of samples:<br>A. Group 3 > Group 1 > Group 2<br>B. Within Group 3, the ranking of samples was based on the following score:<br>$\text{Score}_{\{\text{response}\}} = \text{TMB}_{\{\text{norm}\}} + 2 * \text{PD-L1}_{\{\text{norm}\}}$<br>C. Within Group 1, the ranking of samples was based on the following score:<br>$\text{Score}_{\{\text{response}\}} = \text{TMB}_{\{\text{norm}\}} + \text{PD-L1}_{\{\text{norm}\}}$<br>D. Within Group 2, the ranking of samples was based on the following score:<br>$\text{Score}_{\{\text{response}\}} = \text{TMB}_{\{\text{norm}\}} - \text{PD-L1}_{\{\text{norm}\}}$ |
| **cSysImmunoOnco**<br>**Top-performer in the BOR sub-challenge** | A score of immune response was computed for each patient using EaSleR<sup>41</sup>, which makes use of elastic-net regularized multitask linear regression models trained on TCGA data using quantitative descriptors of the TME as model input and 10 published transcriptomic signatures of immune response as model output. The quantitative descriptors of the TME included relative abundances of different immune cell types,<sup>42</sup> scores of pathway<sup>43</sup> and transcription factor activities,<sup>44</sup> and scores of inter-cellular communication and were derived by combining prior knowledge about the tumor microenvironment and patients' transcriptomics data. The models were fine-tuned by associating penalties with markers of tumor foreignness based on TMB, wherever available, or MSI status estimated using an RNA-seq based signature. |
| **DukeLKB1**<br>**Top-performer in the OS sub-challenge** | A model with six derived features (TMB, PD-L1, 4-gene inflammatory signature, *LKB1* loss signature, *NRF2* activation signature, and neuroendocrine differentiation signature) was generated.<sup>45,46</sup><br>The scores included in the model were calculated as follows: for TMB and PD-L1 components, tumors with respective phenotype > 67<sup>th</sup> percentile were given a score of 1, |
|  | and remaining tumors were scored 0. The 4-gene inflammatory signature and the three tumor-intrinsic gene expression variables were taken as means of the scaled expression scores for the corresponding signature genes. Because we anticipated differences in gene expression and distribution according to tumor histology, the dataset was first separated into squamous and non-squamous subsets, with scaling and averaging across genes performed separately between the two groups. |
| **FICAN-OSCAR**<br>**Top-performer in the OS sub-challenge** | A single linear regression model using a novel **Optimal Subset CArdinality Regression** (*oscar*) L0-quasinorm regularization was generated using the R package available at https://github.com/Syksy/oscar/releases/tag/v0.6.1.<sup>47,48</sup> The model is a linear product of the data matrix X and regularized beta coefficients b. Gene expression signature (CUSTOM FOPANEL) was estimated using a custom gene panel analyzed with GSVA (with the parameter *mx.diff* = TRUE). Other variables included in the model were sex, histology (squamous vs not), smoking history, ECOG performance status (0 vs not), TMB, and PD-L1. A description of each coefficient is available in Supplementary Methods 1.<br>FICAN-OSCAR model equation:<br>$Y = -0.693 \times \text{CUSTOM\_FOPANEL} - 0.357 \times \text{isTMBhigh} - 0.105 \times \text{isMale} - 0.198 \times \text{isSquamous} - 0.05 \times \text{isSquamous\&Above5PDL1} - 0.223 \times \text{isEversmoker} - 0.105 \times \text{isECOG0}$ |
| **@jacob.pfeil**<br>**Top-performer in the OS sub-challenge** | The AbbVie Taux model used an unbiased feature engineering strategy to identify gene expression ratios that differentiate anti-PD-1 responders from non-responders. The reason for using gene expression ratios was to down-weight the effect of response markers by a factor proportional to resistance marker expression level. Cross-validation and regularization were used to mitigate overfitting on the small number of available training samples. An SVM with radial basis function kernel identified a non-linear boundary separating the responder ratio values from non-responder values. Predictive gene expression ratios balanced markers of response (eg, immune cell markers, Type-I interferon, HLA presentation) with markers of resistance (eg, proliferation and inhibitors of immune recognition) |
| **I-MIRACLE**<br>**Top-performer in the OS sub-** | A rule-based prediction model was generated based on classifying TMB and PD-L1 as high or low as follows:<br>• TMB: TMB values were classified as high if greater than or equal to the upper tertile |
| challenge | and as low otherwise. When TMB was missing, the proliferation score<sup>49</sup> was used as a proxy, as it correlates highly with TMB in NSCLC (see prediction of OS sub-challenge)<br>○ The proliferation score was calculated for each patient using the yaGST R package (http://github.com/miccec/yaGST).<sup>50</sup> Patients with missing TMB were classified as TMB high if their proliferation score was greater than or equal to the upper tertile and as TMB low otherwise<br>• PD-L1: Patients were classified as PD-L1 high if their PD-L1 value was $\geq 50$ and PD-L1 low otherwise. When PD-L1 values were missing, the ICR score was used instead<br>○ The ICR score was derived from a 20-gene signature that reflects the presence of a Th1/cytotoxic immune response.<sup>13</sup> The ICR score was calculated for all patients using the yaGST R package. Patients with missing PD-L1 were classified as PD-L1 high if their ICR score was greater than or equal to the upper tertile and as PD-L1 low otherwise<br>• Patients were given a I-MIRACLE score of 1, 2, or 3 based on their TMB and PD-L1 values, as shown in Figure 3B and in Supplementary Methods 1. If TMB was high (or the proliferation score was high when TMB was missing) and PD-L1 expression was high (or the ICR score was high when PD-L1 was missing), we gave a score of 3. A score of 1 was given when both TMB/proliferation score and PD-L1/ICR were low. A score of 2 was given otherwise |
| **Netphar**<br>**Top-performer in the PFS sub-challenge** | A decision tree-based model was generated using TMB high ($\geq 243$) or low ($< 243$) as a first branching point (prior knowledge: TMB is necessary but not sufficient for triggering the checkpoint inhibitor response) and the expression of PD-L1 in the TMB high branch as the second branching point. The model was designed to be conservative on the TMB low branch with all predictions equal to zero.<br>Model equation: $Y = 10 \times \text{TMB}_{\text{binarized}} + \text{TMB}_{\text{binarized}} \times \text{PD-L1}$ |
| **Team TIDE**<br>**Top-performer in the BOR sub-** | The model integrated TIDE<sup>22</sup> with other clinical phenotypes (eg, PD-L1, TMB, and smoking) by the rank aggregation method to enhance the prediction performance on patient survival and response. Treatment-naïve ICI clinical trial data from the TIDE database and late-stage chemotherapy patients of LUAD, LUSC, and SKCM from TCGA |
| **challenge** | were used as the training data. C-index values for survival with each feature within individual cohort and rank features were calculated according to a custom scoring metric.<br>Features such as TMB, PD-L1, CTL, SMOKE, Dysfunction, Exclusion, T.cell.CD4.non.regulatory from QUANTISEQ,<sup>42</sup> B-cell naive from xCell,<sup>51</sup> IFNG signature, and antigen presentation by MHC-I were selected in the model prediction. |

Abbreviations: BOR, best overall response; CTL, cytotoxic T lymphocytes; EaSleR, estimate systems immune response; ECOG, Eastern Cooperative Oncology Group; GSVA, gene set variation analysis; HLA, human leukocyte antigen; ICI, immune checkpoint inhibitor; ICR, immune constant of rejection; IFNG, interferon gamma; LUAD, lung adenocarcinoma; LUSC, lung squamous cell carcinoma; MHC-I, major histocompatibility complex I; MSI, microsatellite instability; NRF2, nuclear factor erythroid 2–related factor 2; NSCLC, non-small cell lung cancer; OS, overall survival; PD-1, programmed death-1; PD-L1, programmed death ligand 1; PFS, progression-free survival; RNA-seq, RNA sequencing; SKCM, skin cutaneous melanoma; SVM, Support Vector Machine; TCGA, The Cancer Genome Atlas; TIDE, tumor immune dysfunction and exclusion; TMB, tumor mutational burden; TME, tumor microenvironment.

### Prediction of Progression-Free Survival

In the PFS sub-challenge, the Netphar and I-MIRACLE models outperformed the TMB baseline model, achieving C-index DSS of 0.19 and 0.087, respectively (Figure 2A). The Netphar model was based on a decision tree positing that high TMB (≥ 243 missense mutations) was necessary but not sufficient to induce a response to nivolumab, and that tumor cell (TC) % PD-L1 expression became relevant only when TMB was high (Figure 2B; Supplementary Methods 1).

**FIG 2.**
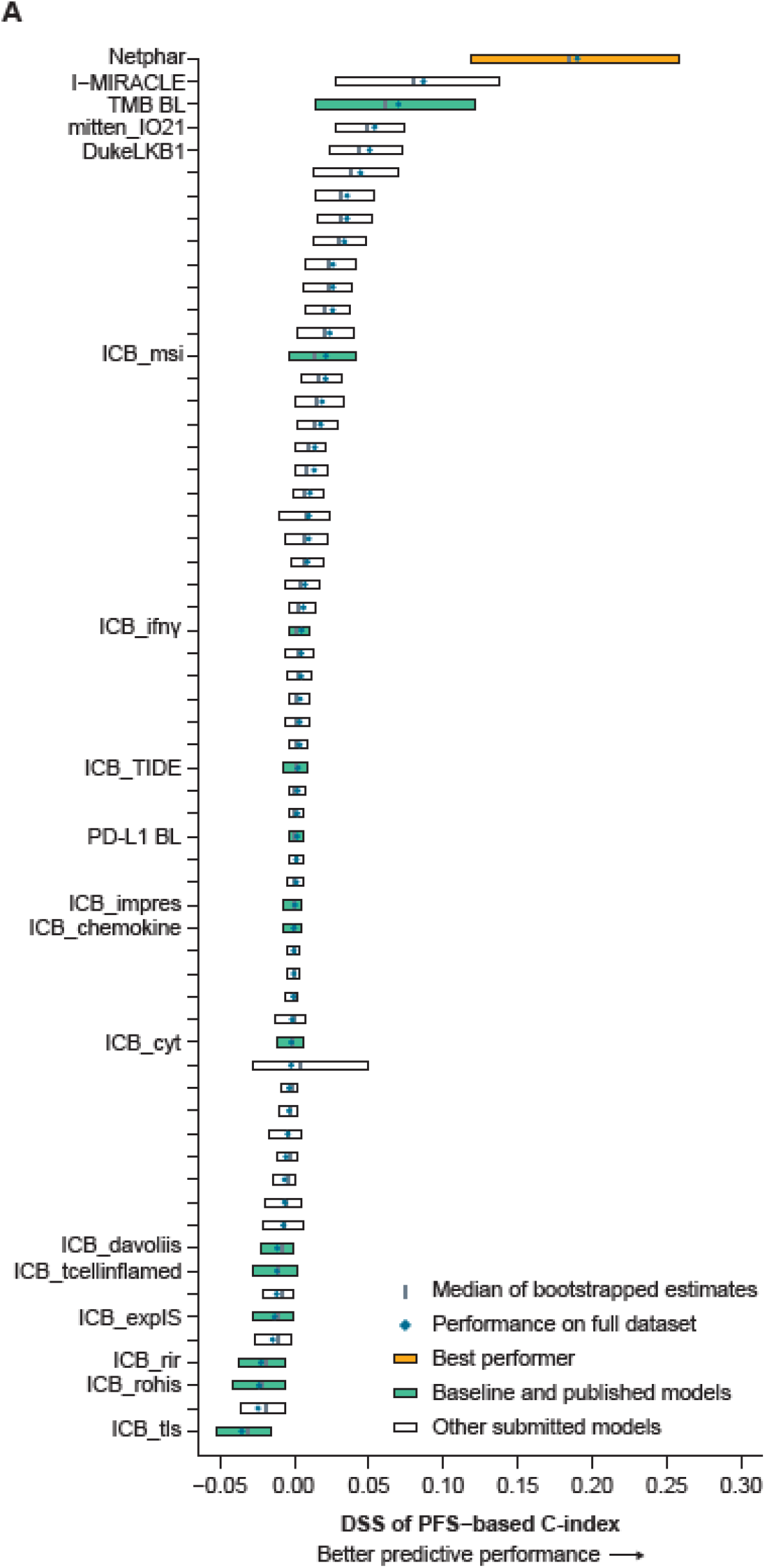
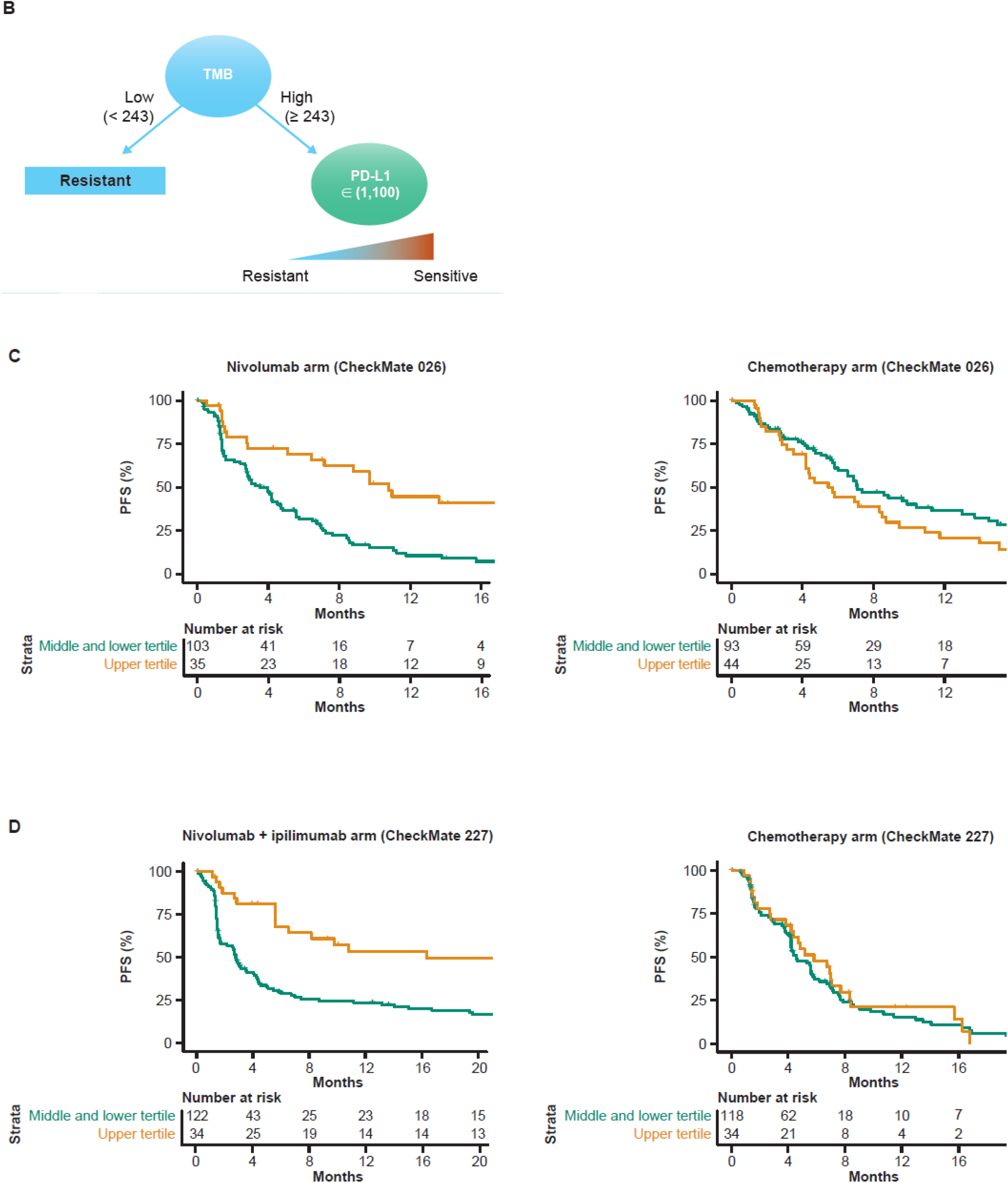
Prediction of PFS with submitted models. (A) Bootstrapped estimates of model performance in CheckMate 026 (boxes are bound by the 25^th^ and 75^th^ percentiles). (B) Decision tree summarizing the Netphar model. (C) Netphar performance in the chemotherapy and nivolumab arms of CheckMate 026. (D) Netphar performance in the chemotherapy and nivolumab + ipilimumab arms of CheckMate 227. BL, baseline; C-index, concordance index; DSS, difference in squared scaled basal metrics; PFS, progression-free survival; PD-L1, programmed death ligand 1; TMB, tumor mutational burden.

In the nivolumab arm of CheckMate 026, patients with Netphar scores in the upper tertile had longer median PFS (10.8 months) than patients with scores in the middle and lower tertiles (3.5 months), whereas in the chemotherapy arm, patients with scores in the middle and lower tertiles had slightly longer median PFS (7.1 months) than patients with scores in the upper tertile (5.4 months) (Figure 2C). Netphar scores in the upper tertile were associated with improved median PFS (16.3 months) in the nivolumab + ipilimumab arm of CheckMate 227 compared with scores in the middle and lower tertiles (2.8 months). In the chemotherapy arm of CheckMate 227, patients with scores in the upper tertile had similar median PFS (5.8 months) to patients with scores in the middle and lower tertiles (4.6 months) (Figure 2D).

### Prediction of Overall Survival

In the OS sub-challenge, three models had higher C-index DSS than baseline models, including TMB and PD-L1, with I-MIRACLE, FICAN-OSCAR, and DukeLKB1 achieving DSS of 0.050, 0.046, and 0.032, respectively (Figure 3A). Although the @jacob.pfeil model had the highest DSS (0.0721), bootstrapped estimates of performance for that model showed substantial variation. The I-MIRACLE model gave patients a score of 1, 2, or 3 based on their TMB and PD-L1 values (Figure 3B and Table 2).

**FIG 3.**
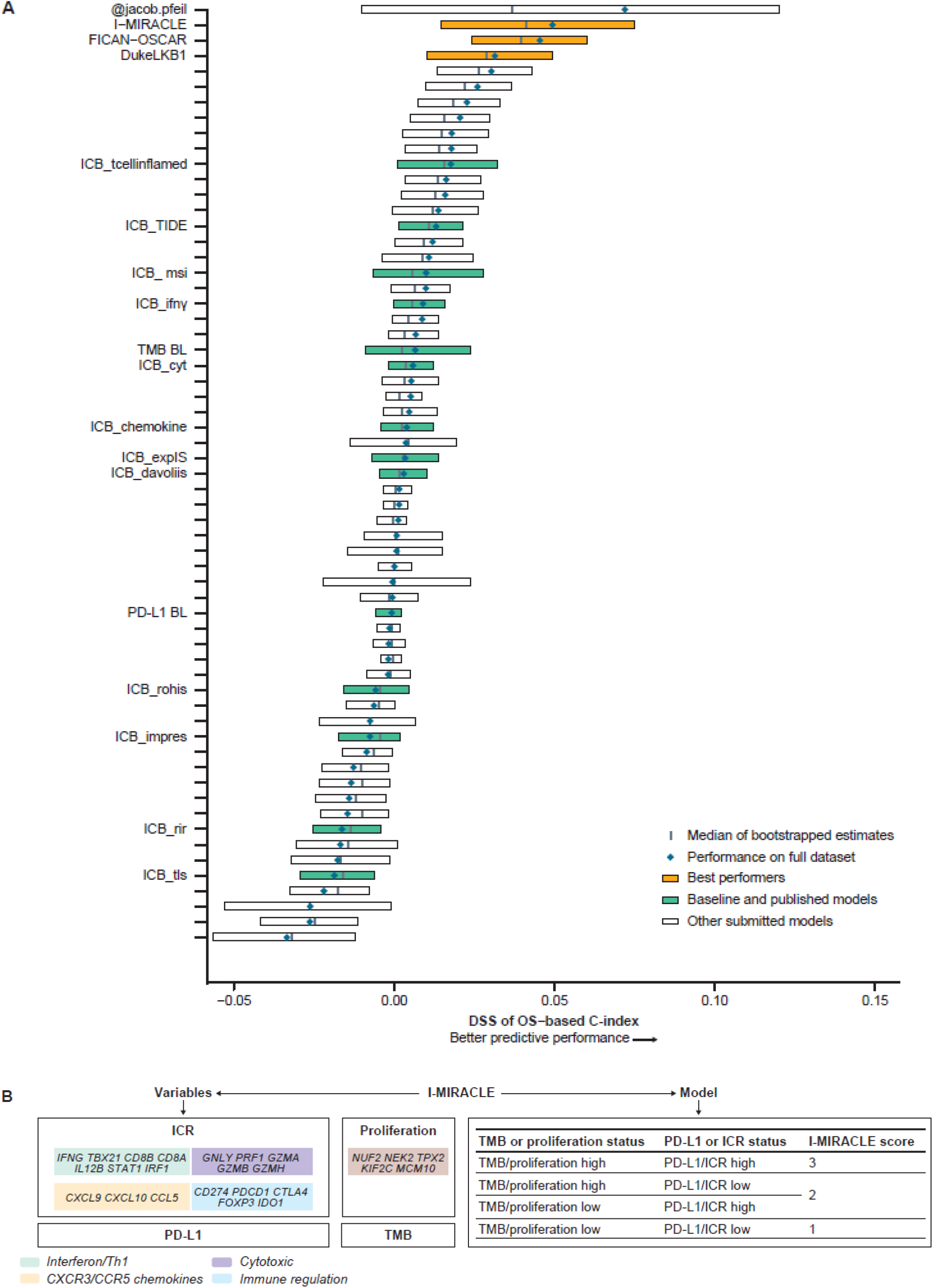
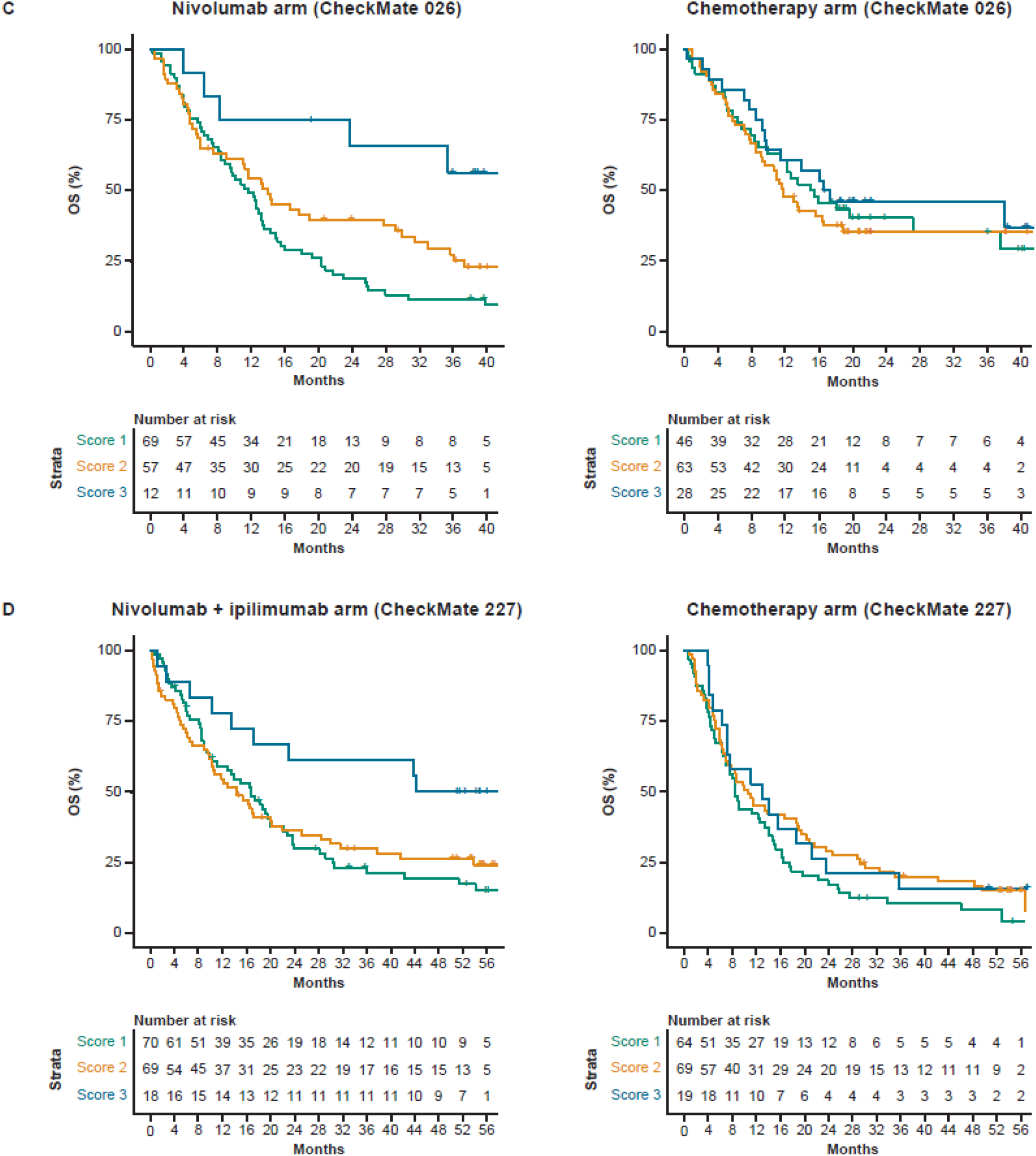
Prediction of OS with submitted models. (A) Bootstrapped estimates of model performance in CheckMate 026 (Boxes are bound by the 25^th^ and 75^th^ percentile). (B) Classification principle of the I-MIRACLE model. (C) I-MIRACLE performance in the chemotherapy and nivolumab arms of CheckMate 026. (D) I-MIRACLE performance in the chemotherapy and nivolumab + ipilimumab arms of CheckMate 227. BL, baseline; C-index, concordance index; DSS, difference in squared scaled basal metrics; ICR, immunologic constant of rejection; OS, overall survival; PD-L1, programmed death ligand 1; TMB, tumor mutational burden.

In the nivolumab arm of CheckMate 026, patients with I-MIRACLE scores of 3 had better median OS (not reached) than patients with scores of 2 (14.1 months) or 1 (11.8 months), whereas in the chemotherapy arm, OS was similar in all patients regardless of I-MIRACLE score (15.2, 11.7, 16.9 months with a score of 1, 2, and 3, respectively) (Figure 3C). In CheckMate 227, I-MIRACLE scores of 3 were associated with prolonged median OS (44.3 months) in the nivolumab + ipilimumab arm compared with scores of 2 (14.3 months) or 1 (16.7 months). OS was similar in the chemotherapy arm regardless of the score (8.5, 10.7, 12.9 months with a score of 1,2, and 3, respectively) (Figure 3D).

### Prediction of Best Overall Response of Progressive Disease

Four models in the BOR sub-challenge surpassed the performance of all baseline models. The DSS of ROC-AUC was 0.055 for cSysImmunoOnco, 0.052 for Aginome-Amoy, 0.049 for Team TIDE, and 0.039 for FICAN-OSCAR (Figure 4A). The cSysImmunoOnco model applied regularized multi-task linear regression to model hallmarks of anticancer immune response based on quantitative descriptors of the tumor microenvironment and TMB (Figure 4B).

**FIG 4.**
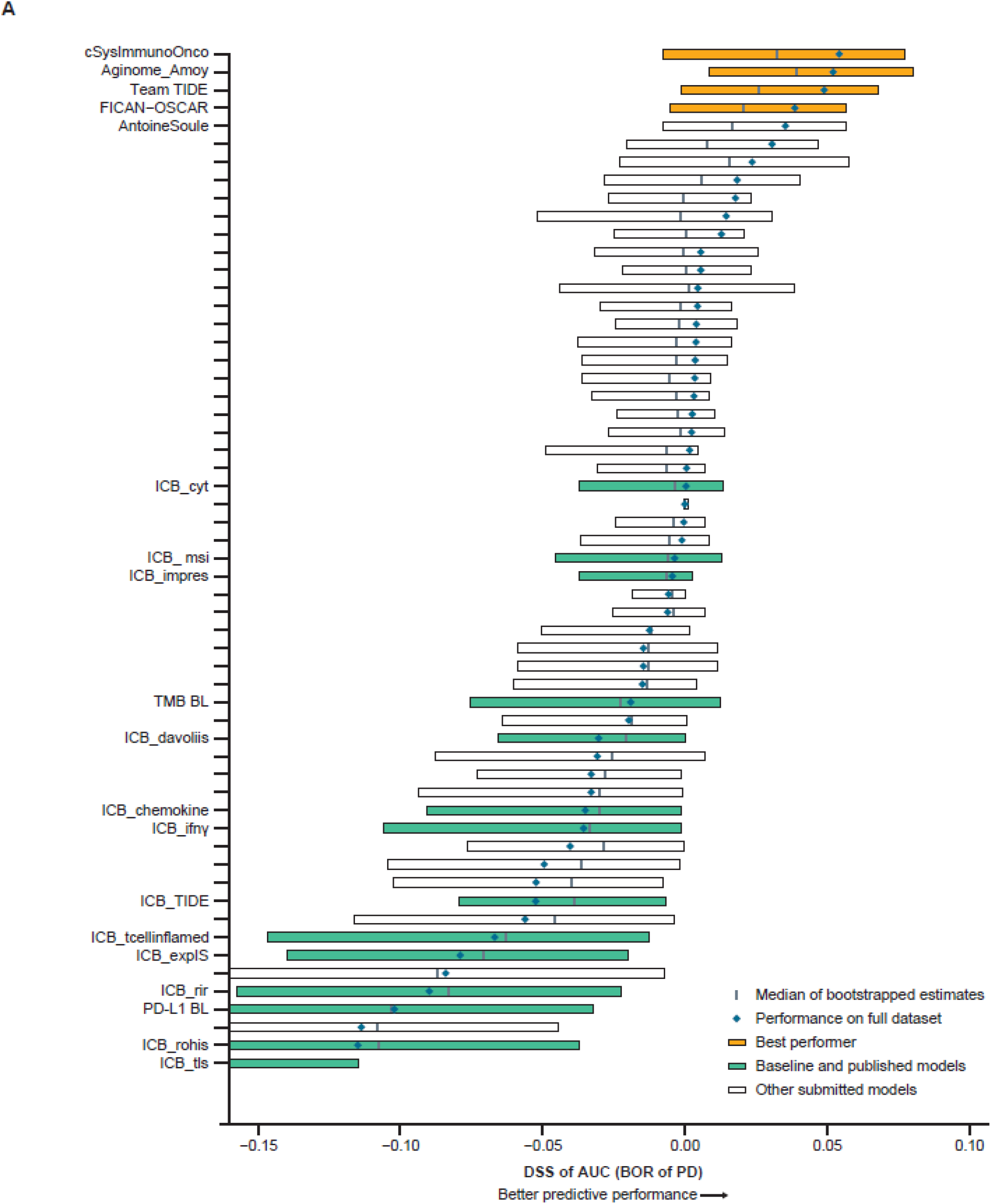
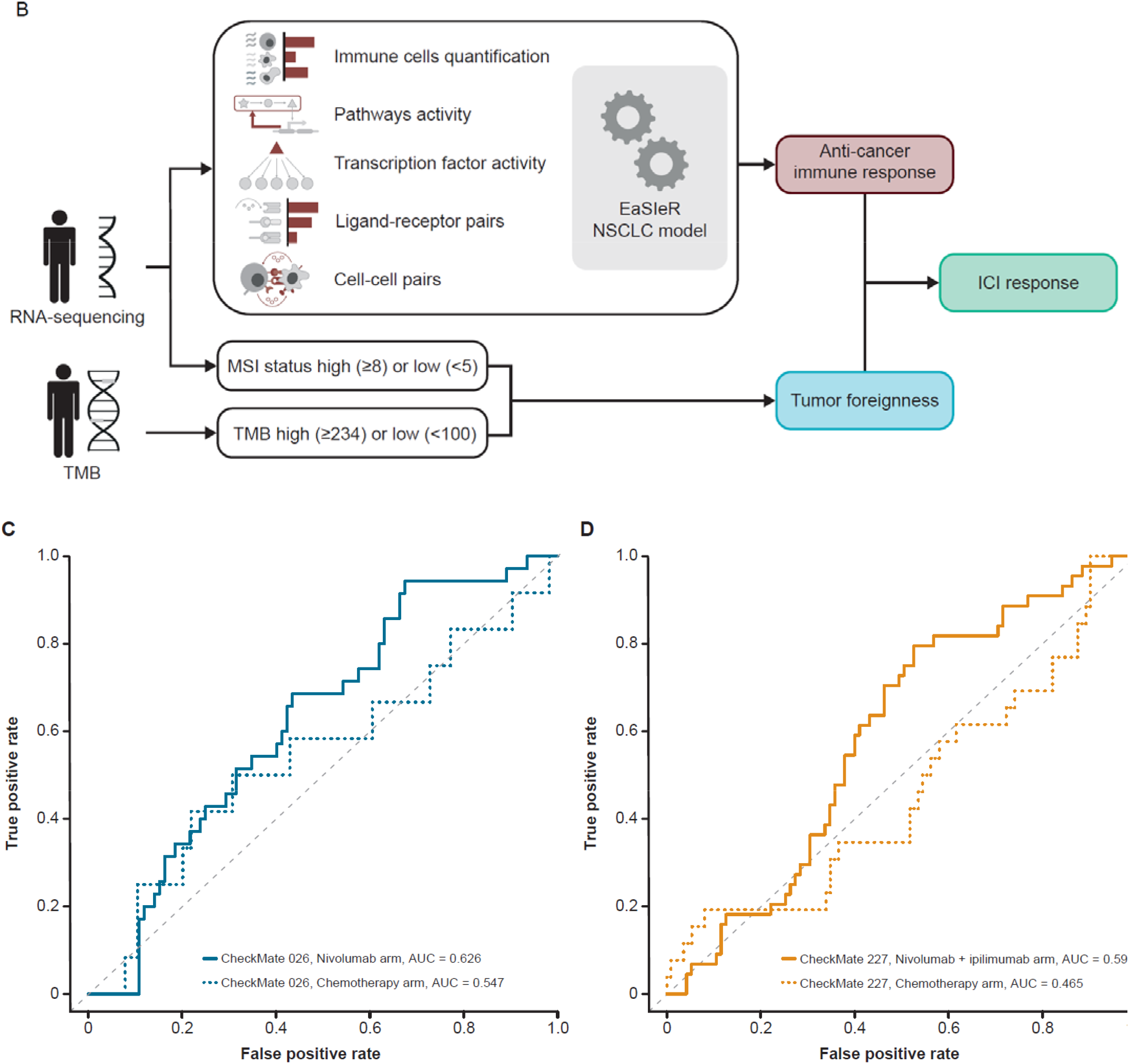
Prediction of BOR of PD with submitted models. (A) Bootstrapped estimates of model performance in CheckMate 026 (boxes are bound by the 25^th^ and 75^th^ percentiles). (B) Principle of the cSysImmunoOnco model. (C) cSysImmunoOnco model performance in CheckMate 026 and (D) CheckMate 227. The grey dotted line is the line of non-determination. AUC, area under the curve; BL, baseline; BOR, best overall response; DSS basal metrics, difference in squared scaled basal metrics; EaSIeR, estimate systems immune response; ICI, immune checkpoint inhibitor; ICR, immunologic constant of rejection; MSI, microsatellite instability; NSCLC, non-small cell lung cancer; OS, overall survival; PD, progressive disease; PD-L1, programmed death ligand 1; TMB, tumor mutational burden.

The ROC-AUC with the cSysImmunoOnco model was higher in the nivolumab arm of CheckMate 026 (0.626) and nivolumab + ipilimumab arm of CheckMate 227 (0.593) than in the chemotherapy arm of CheckMate 026 (0.547) or the chemotherapy arm of CheckMate 227 (0.465) (Figure 4C and 4D).

### Model Performance

Several models had similar or better performance in CheckMate 227 than in CheckMate 026 (Supplementary Figure 2). Netphar was the top-performing model for PFS prediction in the nivolumab arm of CheckMate 026 and in the nivolumab + ipilimumab arm of CheckMate 227. The Netphar model had good predictive accuracy for OS in the nivolumab + ipilimumab arm of CheckMate 227. The I-MIRACLE model had good predictive accuracy for PFS in CheckMate 026 (Supplementary Table 6). The cSysImmunoOnco model did not have good predictive accuracy for PFS or OS in CheckMate 026.

### Gene Signatures

Multiple teams (cSysImmunoOnco, I-MIRACLE, Team TIDE, and FICAN-OSCAR) leveraged publicly available gene expression data to train the models and deemed the expression of a select assortment of genes important (Supplementary Methods 3). The DukeLKB1 six-feature model included a validated transcriptional signature of STK11 functional loss as a predictive feature.^34^ Among the models relying on gene expression information, the cSysImmunoOnco model used the expression of > 100 genes, whereas FICAN-OSCAR relied on five genes (Supplementary Figure 3A). A total of 140 genes ranked important by various models were selected as seeds for downstream analysis. Additional genes that were highly correlated to the seed genes (correlation > 0.85) were included to form a set of 403 genes grouped into three clusters using hierarchical clustering (Supplementary Figure 3B). Analysis of the three clusters showed the enrichment of three main mechanisms. The first cluster represented pathways relevant to tumor intrinsic cell-cycle dysregulation (Supplementary Figure 3Ci, Di). The second cluster included pro-inflammatory immune signatures related to interferon-gamma signaling and antigen presentation (Supplementary Figure 3Cii, Dii). The third cluster included immunosuppressive signatures related to interleukin-10 signaling. The *P* values associated with the third cluster were not small, suggesting weak enrichment, likely due to the small cluster size (Supplementary Figure 3Ciii, Diii). These results show an association of the top predictive genes from the benchmarked models with well-established pathways related to cell-cycle dysregulation and pro-inflammatory immune response.

## Discussion

Studies reporting associations with ICI response in NSCLC have been limited by small sample sizes from single ICI treatment arms.^15,17,18^ This Challenge addressed these shortcomings by using two large and well-characterized phase III RCTs and by comparing predicted responses between ICI- and chemotherapy-treated arms, thereby distinguishing treatment response prediction from prognostic effects. The model-to-data framework was an important characteristic of this Challenge. While participants received limited feedback with this paradigm during model development, which prevented model refinement, this ensured an unbiased and reproducible assessment of the Challenge models.^19^ The model-to-data framework could be made accessible to support evaluation of in silico predictors using various datasets while maintaining data privacy. This study established a robust standard for researchers aiming to identify biomarkers predictive of ICI efficacy. We expect that future Challenges will support efficient biomarker discovery across multiple contexts.

Participants integrated prior knowledge of ICIs with modeling methods like decision trees and regularized regression, additive models with hand-crafted weights, and decision trees with additive models. Preliminary attempts to aggregate models did not show improvements over individual models. While submitted models significantly outperformed TMB and PD-L1 as univariate predictors, most of the top-performing models included both variables, sometimes combined with gene expression signatures such as ICR or a proliferation signature, which reflected the clinical importance of TMB and PD-L1. This aligns with the observations obtained in tumor types, including head and neck squamous cell carcinoma (HNSCC) and melanoma, in which a T-cell–inflamed gene expression profile (similar to ICR) and TMB predicted PFS in patients receiving pembrolizumab. Likewise, a combined assessment of TMB and an inflammatory signature predicted BOR, PFS, and OS in patients with advanced melanoma receiving nivolumab or nivolumab + ipilimumab.^35^ A high ICR score predicted survival or response in patients with multiple tumor types treated with ICIs.^13^

These results indicate that a combination of PD-L1, TMB, and immune gene signatures might be able to identify a subgroup of patients with NSCLC likely to respond to ICI and could be used for the design of a prospective phase III trial or to guide treatment choice. There is no single ‘magic bullet’ biomarker or model-building approach to predict response to ICIs. The biomarker content of top-performing models, as well as the exploration of their gene signature content, reinforce the need to assess tumor biology, tumor immunogenicity, and immune system status to identify patients most likely to benefit from ICI treatment. However, top-performing models differed across subchallenges, suggesting that composite models have different predictive potential, depending on the clinical end point assessed. For example, TMB and PD-L1 seem important for the prediction of PFS and OS, confirming previous studies,^36^ while mechanisms such as apoptosis, T-cell cross talk, and adaptive immune resistance seem important for the prediction of response. Future precision medicine approaches will benefit from the exploration and development of targeted composite biomarker strategies.

The models identified may be generalizable to ICI datasets other than first-line treatment in metastatic NSCLC. Contributing teams used training datasets from other tumor types (melanoma or HNSCC), and the top-performing models in CheckMate 026 were validated in CheckMate 227 with different primary end points. These observations suggest that this approach may provide a blueprint to support modeling initiatives in diverse tumor types.

A possible limitation of this study is that TMB, frequently used in the submitted models, may be inferred from DNA or RNA sequencing data and is a proxy for tumor ‘foreignness’ but does not capture neoantigen clonality and abundance or non-canonical neoantigens generated from other tumor aberrations.^37,38^ Data such as T-cell/B-cell receptor repertoire, tobacco use, ECOG PS, age, and sex are not readily available in public datasets, participants did not always use them, and their role in predicting response to ICIs needs to be explored further. NSCLC is a genetically heterogeneous disease^39^, and specific subpopulations may differ in optimal biomarkers predictive of therapy response. While transcriptional signatures predictive of functional STK11 and KEAP1/NFE2L2 alterations were used in some models, integration of transcriptional phenotypes with fuller exome datasets across larger cohorts will be necessary to discover these subtype-specific biomarkers. Other limitations were the similarity of PFS and OS between the nivolumab and chemotherapy treatment groups of CheckMate 026, and the exclusion of patients with PD-L1 expression < 1% in CheckMate 026. Although clinical and molecular data sets from both trials are large and rich, ascertainment of genomics data was incomplete because of logistical limitations. When the CheckMate 026 and 227 studies were conducted, chemotherapy was the standard of care; the current standard is chemotherapy plus ICI.^40^ The models identified here should be tested in the context of this new standard.

This pioneering study showed that a crowdsourced approach could successfully identify clinical and translational characteristics predictive of ICI efficacy. This analysis improves the understanding of the mechanisms of tumor sensitivity and resistance to treatment, which will support the development of therapies for patient subpopulations unlikely to benefit from current ICI regimens. It provides a roadmap for successful partnership between academic and industry scientists that allows for robust, reproducible biomarker testing while protecting patient data and incentivizing collaboration. We hope that the DREAM Challenge framework will be used to analyze data from many phase III trials, to speed the development of clinically actionable biomarkers and improve patient outcomes.

## Acknowledgments

Teemu D. Laajala was funded by the Finnish Cancer Institute and the Finnish Cultural foundation as a FICAN Cancer Researcher. Anni S. Halkola received funding from the University of Turku Graduate School (MATTI), the Academy of Finland (grants 310507, 313267 and 326238), the Cancer Society of Finland, and the Sigrid Jusélius Foundation. Mikko I. Mäyränpää received funding from the Finnish Medical Foundation. Tuomas Mirtti received funding from the Academy of Finland.

Francesca Finotello was supported by the Austrian Science Fund (FWF) [T 974-B30] and by the Oesterreichische Nationalbank (OeNB) [18496]. Óscar Lapuente-Santana was supported by the Department of Biomedical Engineering, Eindhoven University of Technology.

Wenyu Wang, Jie Bao, and Jing Tang were supported by an ERC Starting Grant (DrugComb, No. 716063), the Academy of Finland (No. 317680), and the Sigrid Jusélius Foundation. Wenyu Wang holds a funded position at the Doctoral program of Biomedicine, University of Helsinki and holds a personal grant from K. Albin Johanssons Stiftelse and Ida Montinin Säätiö. Emmy Verschuren was supported by the Academy of Finland (No. 328437), the iCAN Digital Precision Cancer Medicine Flagship (No. 320185 Academy of Finland), and the CAN-PRO Translational Cancer Medicine Research Program Unit. Data analysis resources were provided by the CSC – IT Center for Science, Finland.

Jacob Kaufman received funding from the Department of Defense (Lung Cancer Research Program Concept Award LC180633) and was the recipient of a SITC-AstraZeneca Lung Cancer Clinical Fellowship (SPS256666).

Lin Yan and Yang Liu received PACT funding through FNIH. Ming Tang received funding from the NIH. Shengqing Stan Gu was the recipient of the Sara Elizabeth O’Brien Trust Fellowship. Avinash Das Sahu received funding from the NCI (K99CA248953) and the Human Immunome Project (MP19-02-190).

Davide Bedognetti received the following grant from Sidra Medicine Internal Funds (SDR400123). Michele Ceccarelli received the following grant AIRC: IG 2018 ID 21846. Xiaole Shirley Liu contributed to the development of the TIDE model. Josue Samayoa contributed to the development of the @jacob.pfeil model. Abraham Apfel contributed data analysis advice.

The study was supported by Bristol Myers Squibb.

Medical writing and editorial support were provided by Thierry Deltheil, PhD, and

Matthew Weddig of Spark Medica Inc., funded by Bristol Myers Squibb.

## Author contributions

## Data sharing statement

More information on Bristol Myers Squibb’s data sharing policy can be found here: https://www.bms.com/researchers-and-partners/clinical-trials-and-research/disclosure-commitment.html

