## Supplement for "A Community Challenge to Predict Clinical Outcomes After Immune Checkpoint Blockade in Non-Small Cell Lung Cancer"

**Supplementary information**

### Author teams’ contributions

If the author contributed to the generation of a model, the name of the model is indicated in parentheses next to their name in the paragraph below. The table below the paragraph shows the ranking of each model for each sub-challenge.

Mike Mason^1^; Óscar Lapuente-Santana (cSysImmunoOnco)^2†^; Anni S. Halkola (FICAN-OSCAR)^3†^; Wenyu Wang (Netphar)^4†^; Raghvendra Mall (I-MIRACLE)^5,6†^; Xu Xiao (Aginome-Amoy)^7,8†^; Jacob Kaufman (DUKELKB1)^9,10†^; Jingxin Fu (Team TIDE)^11†^; Jacob Pfeil (@jacob.pfeil)^12†^; Jineta Banerjee^13^; Verena Chung^13^; Han Chang^1^; Scott D. Chasalow^1^; Hung Ying Lin^1^; Rongrong Chai^13^; Thomas Yu^13^; Francesca Finotello (cSysImmunoOnco)^14,15^; Tuomas Mirtti (FICAN-OSCAR)^16,17,18,19^; Mikko I. Mäyränpää (FICAN-OSCAR)^16^; Jie Bao (Netphar)^4^; Emmy W. Verschuren (Netphar)^20^; Eiman I. Ahmed (I-MIRACLE)^21^; Michele Ceccarelli (I-MIRACLE)^22,23^; Lance D. Miller (I-MIRACLE)^24,25^; Gianni Monaco (I-MIRACLE)^23^; Wouter R.L. Hendrickx (I-MIRACLE)^21,26^; Shimaa Sherif (I-MIRACLE)^21,26^; Lin Yang (Team TIDE)^11^; Ming Tang (Team TIDE)^11^; Shengqing Stan Gu (Team TIDE)^11^; Wubing Zhang (Team TIDE)^11^; Yi Zhang (Team TIDE)^11^; Zexian Zeng (Team TIDE)^11^; Avinash Das Sahu (Team TIDE)^11^; Yang Liu (Team TIDE)^11‡^; Wenxian Yang (Aginome-Amoy)^27‡^; Davide Bedognetti (I-MIRACLE)^21,26,28‡^; Jing Tang (Netphar)^29‡^; Federica Eduati (cSysImmunoOnco)^2,30‡^; Teemu D. Laajala (FICAN-OSCAR)^3,31,32‡^; William J. Geese^1^; Justin Guinney^33^*; Joseph D. Szustakowski^1^*; David P. Carbone^10^*; Benjamin G. Vincent^34^*

**Authors’ affiliations:**

^1^Bristol Myers Squibb, Princeton, NJ, USA

^2^Department of Biomedical Engineering, Eindhoven University of Technology, Eindhoven, The Netherlands

^3^Department of Mathematics and Statistics, University of Turku, Turku, Finland

^4^Faculty of Medicine, University of Helsinki, Helsinki, Finland

^5^Qatar Computing Research Institute, Hamad Bin Khalifa University, P.O. Box 34110, Doha, Qatar

^6^Department of Immunology, St. Jude Children's Research Hospital, P.O. Box 38105, Memphis, TN, USA

^7^School of Informatics, Xiamen University, Xiamen, China

^8^National Institute for Data Science in Health and Medicine, Xiamen University, Xiamen, China

^9^Department of Medicine, Duke University, Durham, NC, USA

^10^The Ohio State University Comprehensive Cancer Center, Columbus, OH, USA

^11^Dana-Farber Cancer Institute, Boston, MA, USA

^12^AbbVie, South San Francisco, CA, USA

^13^Sage Bionetworks, Seattle, WA, USA

^14^Institute of Molecular Biology, University of Innsbruck, Innsbruck, Austria

^15^Digital Science Center (DiSC), University of Innsbruck, Innsbruck, Austria

^16^Department of Pathology, University of Helsinki and Helsinki University Hospital, Helsinki, Finland

^17^Research Program in Systems Oncology, University of Helsinki, Helsinki, Finland

^18^iCAN-Digital Precision Cancer Medicine Flagship, Helsinki, Finland

^19^Department of Biomedical Engineering, School of Medicine, Emory University, Atlanta, GA, USA

^20^Institute for Molecular Medicine Finland (FIMM), HiLIFE, University of Helsinki, Helsinki, Finland

^21^Human Immunology Department, Sidra Medicine, P.O. Box 26999, Doha, Qatar

^22^Department of Electrical Engineering and Information Technology (DIETI), University of Naples "Federico II", 80125 Naples, Italy

^23^BIOGEM Institute of Molecular Biology and Genetics, Via Camporeale, Ariano Irpino, Italy

^24^Department of Cancer Biology, Wake Forest School of Medicine, Winston-Salem, NC, USA

^25^Atrium Health Wake Forest Baptist Comprehensive Cancer Center, Winston-Salem, NC, USA

^26^College of Health and Life Sciences, Hamad Bin Khalifa University, P.O. Box 26999, Doha, Qatar

^27^Aginome Scientific, Xiamen, China

^28^Department of Internal Medicine and Medical Specialties, University of Genoa, Genoa, Italy

^29^University of Helsinki, Helsinki, Finland

^30^Institute for Complex Molecular Systems (ICMS), Eindhoven University of Technology, Eindhoven, The Netherlands

^31^FICAN West Cancer Centre, University of Turku and Turku University Hospital, Turku, Finland

^32^Department of Pharmacology, Anschutz Medical Campus, University of Colorado, Denver, CO, USA

^33^Tempus Labs, Chicago, IL, USA

^34^Department of Medicine, Division of Hematology, Department of Microbiology and Immunology, Curriculum in Bioinformatics and Computational Biology, Computational Medicine Program, University of North Carolina at Chapel Hill, Chapel Hill, NC, USA

^†^Lead authors from participating teams in the Anti–PD-1 Response Prediction DREAM Challenge with equal contribution

^‡^Senior authors from participating teams in the Anti–PD-1 Response Prediction DREAM Challenge with equal contribution

*Co-senior authors from the Anti–PD-1 Response Prediction DREAM Challenge steering committee

**Summary of** **top-performing models and breakdown of each participating team and ranking of models**

|  | **PFS sub-challenge** | **OS sub-challenge** | **BOR sub-challenge** |
| --- | --- | --- | --- |
| **Summary of top-performing models by sub-challenge^a^** | Netphar | I-MIRACLE  FICAN-OSCAR  DukeLKB1  @jacob.pfeil^b^ | Aginome_Amoy  cSysImmunoOnco  Team TIDE  FICAN-OSCAR |
| **Model/team name and participants^c^** | **Model ranking^a^** | | |
| Aginome_Amoy   - Xu Xiao - Wenxian Yang | 11 | 16 | 1^c^ |
| cSysImmunoOnco team   - Óscar Lapuente-Santana - Francesca Finotello - Federica Eduati | 6 | 25 | 1^c^ |
| DukeLKB1   - Jacob Kaufman | 4 | 1^c^ | 23 |
| FICAN-OSCAR   - Anni S. Halkola - Tuomas Mirtti - Mikko I. Mäyränpää - Teemu D. Laajala | 9 | 1^c^ | 1^c^ |
| I-MIRACLE   - Raghvendra Mall - Eiman I. Ahmed - Michele Ceccarelli - Lance D. Miller - Gianni Monaco - Wouter R. L. Hendrickx - Shimaa Sherif - Davide Bedognetti | 2 | 1^c^ | 9 |
| @jacob.pfeil   - Jacob Pfeil | 22 | 1^b^ | 16 |
| Netphar   - Wenyu Wang - Jie Bao - Emmy W. Verschuren - Jing Tang | 1 | 19 | 22 |
| Team TIDE   - Jingxin Fu - Lin Yang - Ming Tang - Shengqing Stan Gu - Wubing Zhang - Yi Zhang - Zexian Zeng - Avinash Das Sahu - Yang Liu | 12 | 8 | 1^c^ |

BOR, best overall response; OS, overall response; PFS, progression-free survival.

^a^Only top-performing models in one of the three sub-challenges are indicated in the table. ^b^Although the @jacob.pfeil model had the highest DSS (0.0721), bootstrapped estimates of model performance showed substantial variation; ^c^indicates statistical ties. ^c^Participants are listed by lead, middle, and senior authors for each team.

### Supplementary Methods 1. Methods used for the development of best-performing models

#### Aginome_Amoy model

The full description of the model is available [here](https://www.synapse.org/#!Synapse:syn24893030/wiki/609056).

1. Features selection

To select the critical features for predicting clinical response, we used the Cox proportional-hazards model on publicly available lung cancer and melanoma datasets to identify the clinical measurements that have the highest correlation with the outcome of anti–programmed death-1 (PD-1) therapies measured by overall survival (OS) and progression-free survival (PFS). Both tumor mutational burden (TMB) and programmed death ligand 1 (PD-L1) scores showed strong correlation with the outcome of anti–PD-1 therapy.

2. Data pre-processing and normalization

We filled the missing values in both TMB and PD-L1 scores with zero, and then we applied min-max normalization among all samples to make each selected feature comparable.

3. Ranking of samples

We divided patients into three non-overlapping groups based on their PD-L1 and TMB values:

- Group 1: PD-L1 score below median
- Group 2: PD-L1 score above median and TMB expression below median
- Group 3: both PD-L1 and TMB expression above median

We observed from published clinical trial results that patients from Group 3 responded most favorably to anti–PD-1 therapy among all these groups, patients from Group 2 responded least favorably, and patients from Group 1 were somewhere in between these two groups. Moreover, we noticed that these two biomarkers seem to exhibit non-additive effects in different patient groups. For example, for patients with PD-L1 scores above median, PD-L1 was a strong positive biomarker only in patients with high TMB. Otherwise, PD-L1 score seemed to correlate more with favorable outcome from chemotherapy rather than anti–PD-1 therapy for Group 2 patients who responded least favorably to anti–PD-1. Unfortunately, due to lack of training data, we were not able to systematically train a suitable model to capture these relationships. Instead, we used the following heuristic rules to decide the ranking of samples:

- Group 3 > Group 1 > Group 2
- Within Group 3, the ranking of samples was based on the following score:
   Score_response_ = TMB_norm_ + 2 * PD-L1_norm_
- Within Group 1, the ranking of samples was based on the following score: 
  Score_response_ = TMB_norm_ + PD-L1_norm_
- Within Group 2, the ranking of samples was based on the following score: 
  Score_response_ = TMB_norm_ – PD-L1_norm_

#### cSysImmunoOnco model

The full description of the model is available [here](https://www.synapse.org/#!Synapse:syn23682259/wiki/608719).

1. Quantitative descriptors of the tumor microenvironment

Following the approach described in Lapuente-Santana et al., 2021,^1^ we combined RNA sequencing (RNA-seq) data with prior knowledge to derive five types of quantitative descriptors of the tumor microenvironment, including composition of the tumor immune contexture and activity of intra- and extra-cellular communication (see Table below). As shown in the table, each descriptor provides quantification of a number of features.

**Derivation of the quantitative descriptors of the tumor microenvironment.**

| **Quantitative descriptors** | **Descriptor conception** | **Prior knowledge** | **Number of features** |
| --- | --- | --- | --- |
| Pathway activity | Activity of signaling pathways was inferred using PROGENy, which applies the weighted sum of the expression of pathway signature genes^2^ | Pathway perturbation-response gene expression signatures^2^ | 14 |
| Immune cell quantification | Cell fractions were derived using quanTIseq, which performs in silico deconvolution of complex tissues profiled with bulk transcriptomics^3^ | Cell-type specific gene expression signatures^3^ | 11 |
| Transcription factor activity | Activity of transcription factors was estimated using DoRothEA, which scores each transcription factor based on the average ranks of each transcription factor target genes^4^ | Networks of transcription factor-target interactions (regulons)^4^ | 118 |
| Ligand-receptor pairs | Weights for ligand-receptor pairs were computed as the minimum of the expression of the ligand and the receptor^1^ | Annotations of ligand-receptor pairs^5^ | 813 |
| Cell-cell interaction | Cell-cell interaction scores were computed as the weighted sum of the number of expressed ligand-receptor pairs. The weights correspond to the inverse of the ligand-receptor pair frequency across the whole of TCGA (hypothesizing that rare interactions are more relevant to discriminate patients)^1^ | Annotations of cell-type specific ligand-receptor pairs^5^ | 169 |

TGCA, The Cancer Genome Atlas.

2. Hallmarks of immune response

We used ten published transcriptomics signatures as hallmarks of anticancer immune response, which we considered as output variables (hereafter called “tasks”). More detailed information can be found in the original work^1^ and in Supplementary Tables 2 and 3.

| **Hallmark of the immune response** | **Original study** |
| --- | --- |
| Cytolytic activity (ICB_cyt) | Rooney et al., *Cell*, 2015^6^ |
| Roh immune score (ICB_rohis) | Roh et al., *Sci Transl Med*, 2017^7^ |
| Chemokine signature (ICB_chemokine) | Messina et al., *Nat Sci Rep*, 2012^8^ |
| Davoli immune signature (ICB_davoliis) | Davoli et al., *Science*, 2017^9^ |
| IFNy signature (ICB_ifnγ) | Ayers et al., *J Clin Invest*, 2017^10^ |
| Expanded immune signature (ICB_expIS) | Ayers et al., *J Clin Invest*, 2017^10^ |
| T-cell inflamed signature (ICB_tcellinflamed) | Ayers et al., *J Clin Invest*, 2017^10^ |
| Repressed immune resistance (ICB_rir) | Jerby-Arnon et al., *Cell*, 2018^11^ |
| Tertiary lymphoid structures signature (ICB_tls) | Cabrita et al., *Nature*, 2020^12^ |

3. Machine learning approach

The objective function that defines the regularized multi-task linear regression for *n* observations is formulated as:

$$\frac{1}{N}\sum_{i=1}^{N} {|\left| yi-\beta0-xi\beta\right||}_{2}^{2} + \lambda\sum_{j=1}^{p} ((1-\alpha\beta_{j}\left| \left| 2+\alpha\right| \right|\beta_{j}{||}_{2}^{2}$$

where $yi$ represents a row-vector of response variables (tasks), and $xi$ is a row-vector of observed features.

The aim of the regularized multi-task linear regression is to estimate a matrix $\beta$ whose rows represent the relation between one feature and all the tasks, and a vector $\beta0$ of offsets (one for each task). The regularization term is a grouped version of the elastic net that aims at enforcing sparsity to entire rows of $\beta$.^13,14^ In this way, the features corresponding to those rows of $\beta$ that are set to zero do not contribute to the model. The strength of the regularization effect is tuned via the hyperparameter $\lambda$, while $\alpha$ regulates the interplay between the ridge- and the lasso-like terms of the elastic net. We selected the hyperparameters using 5-fold cross-validation. We used regularized multi-task linear regression as implemented in the glmnet R package 2.0-16.^15^

4. Model training

Models were trained on a combined RNA-seq dataset from The Cancer Genome Atlas (TCGA) patients with lung adenocarcinoma (LUAD) or lung squamous cell carcinoma (LUSC); only data from primary tumors were considered. The model training was repeated 100 times with random partitioning, each time picking 20% of the samples as a test set. This helped to assess the stability of the model, both in terms of performance and feature selection. For each iteration, we first standardized the training set, and then we standardized the test set based on the mean and standard deviation of the training set.

5. Prediction of response

For each of the five quantitative descriptors (eg, activity of the 14 features associated with pathway activity), all the models identified in the randomized cross-validation (100 runs) were used to make predictions for the patients from this Challenge dataset. Given the multitask problem, each of these models was in fact a composition of 10 models, one for each task. Therefore, predictions for each patient were then computed as the average across runs, tasks, and individual quantitative descriptors.

Using our TCGA-trained models, we could predict immune response, which is required for effective immunotherapy. To additionally account for tumor foreignness, another hallmark of effective immunotherapy, we introduced an additional penalty based on TMB. We first selected patients with high TMB (TMB ≥ 234) and low TMB (TMB < 100), as described in Hellmann et al., 2018.^16^ For patients where TMB was not available, we used a transcriptomic signature quantifying microsatellite instability (MSI) status^17^ to score them as MSI high (MSI ≥ 8) or MSI low (MSI < 5), based on the relationship between TMB and MSI assessed on the TCGA data.

To predict best overall response (BOR), we then refined our predictions by adding a positive penalty to TMB (MSI)-high and a negative penalty to TMB (MSI)-low patients. The penalty values were empirically defined based on model predictions on melanoma patients treated with anti–PD-1 therapy^18^ and on leaderboard results.

For the final predictions, TMB (MSI)-high patients’ predicted values were increased by 0.7, whereas TMB (MSI)-low patients were reduced by -0.3. We applied min-max normalization to provide values between 0 and 1.

The strongest predictive biomarkers included those associated with apoptosis (Trail pathway) and T-cell crosstalk (interaction between CD8+ T cells and dendritic cells), as well as markers of adaptive immune resistance (JAK-STAT pathway and STAT4 transcription factor). In the figure below, bar plots show the estimated weights for the top 20 biomarkers. The median of the weights computed first across 100 iterations and then across tasks are shown. Biomarkers are sorted by the absolute mean of their median weights.

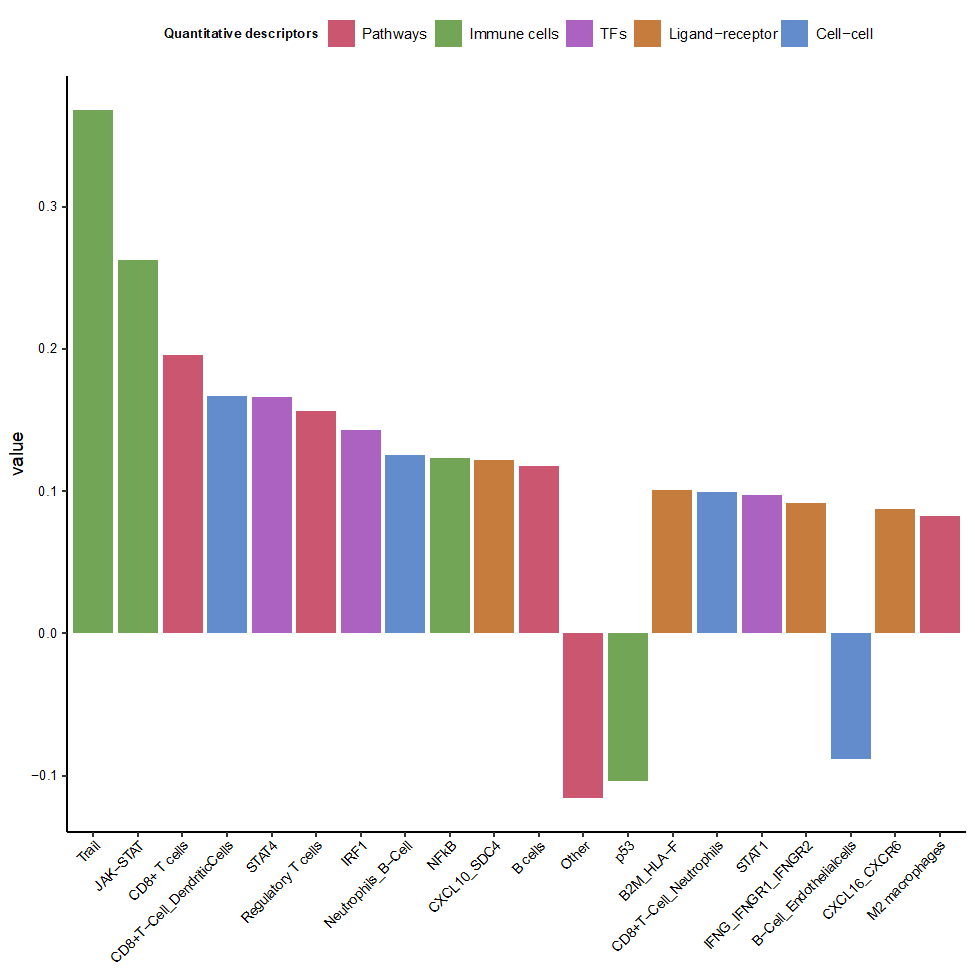

Biomarker weight

The code used for building the model is available at <https://github.com/olapuentesantana/mechanistic_biomarkers_immuno-oncology>.

The EaSIeR package to use RNA-seq data to predict immune response based on quantitative descriptors of the tumor microenvironment (part of the source code submitted through Synapse) is also available as an R/Bioconducor package (<https://www.bioconductor.org/packages/release/bioc/html/easier.html>).

#### DukeLKB1 model

The full description of the model is available [here](https://www.synapse.org/#!Synapse:syn23662130/wiki/609003).

We calculated six tumor variables to construct our models. Three were combined to calculate a baseline model that was expected to have positive association with nivolumab outcomes in the CheckMate 026 study and included PD-L1 expression, TMB, and the 4-gene inflammatory signature (*CD274*, *CD8A*, *LAG3*, *STAT1*).^19^ The remaining three variables represented
LKB1-loss, defined by a 16-gene signature,^20^ NRF2 activation, defined by a 19-gene expression signature (unpublished), and neuroendocrine differentiation, defined by a 22-gene expression signature (unpublished). Based on our preliminary data, we predicted that each of these three tumor-intrinsic phenotypes would be associated with resistance to nivolumab. Given the specified design of the DREAM Challenge, allowing for a total of five model submissions, we combined these variables in the following models, which were submitted to each of the three sub-challenges. Submission ID for PFS sub-challenge and performance characteristics are shown in the table below.

| **Model  (submission ID)** | **Model components** | **Primary metric [bootstrapped estimates]** | **C-index of PFS in the nivolumab arm** | **C-index of PFS in the chemotherapy arm** |
| --- | --- | --- | --- | --- |
| Model 1: Baseline (9710334) | TMB + PD-L1 + 4-gene inflammatory signature | 0.0371  [0.012, 0.0532] | 0.6033 | 0.4782 |
| Model 2: Intrinsic LKB1/NRF2 (9710337) | -LKB1 - NRF2 | -0.1558 [‑0.2013, ‑0.1258] | 0.5589 | 0.7064 |
| Model 3: Combined LKB1/NRF2 (9710340) | TMB + PD-L1 + 4-gene inflammatory signature - LKB1 - NRF2 | 0.0329  [0.0131, 0.0437] | 0.6005 | 0.5351 |
| Model 4: Intrinsic LKB1/neuroendocrine differentiation (9710343) | -LKB1 - neuroendocrine differentiation | -0.1749 [‑0.2954, ‑0.026] | 0.5978 | 0.7273 |
| Model 5: Combined LKB1/neuroendocrine differentiation (9710346) | TMB + PD-L1 + 4-gene inflammatory signature - LKB1 - neuroendocrine differentiation | 0.0556  [0.0367, 0.0792] | 0.622 | 0.4949 |

The scores included in the models were calculated as follows: for TMB and PD-L1 components, tumors with respective phenotype greater than the 67^th^ percentile were given a score of 1, and tumors at or below the 67^th^ percentile were scored 0. The 4-gene inflammatory signature and the three tumor-intrinsic gene expression variables were taken as means of the scaled expression scores for the corresponding signature genes. Because we anticipated differences in gene expression and distribution according to tumor histology, the dataset was first separated into squamous and non-squamous subsets, with scaling and averaging across genes performed separately between the two groups.

| **Signature** | **Signature genes** | **Variables by histology** | **Algorithm** |
| --- | --- | --- | --- |
| 4-gene inflammatory signature score^19^ | *CD274, CD8A, LAG3, STAT1*  Normalization: TMM (NOISeq) | inflam_sig SQ: Median of scaled expression for four genes across squamous tumors  inflam_sig NSQ: Median of scaled expression for four genes across non-squamous tumors | Varinflam: If inflam_sig > 0.5, Varinflam = 0.5; if inflam_sig < -0.5, varinflam = -0.5; Else, varinflam = 0 |
| PD-L1 | **-** | **-** | VarPD-L1: If PD-L1 > 67^th^ percentile, then VarPD-L1 = 1; Else Var$PD-L1 = 0 |
| TMB | **-** | **-** | VarTMB: IfTMB >67^th^ percentile, then VarTMB =1; Else Var$TMB = 0 |
| LKB1 loss score^20^ | *AVPI1, BAG1, CPS1, DUSP4, FGA, GLCE, HAL, IRS2, MUC5AC, PDE4D, PTP4A1, RFK, SIK1, TACC2, TESC, TFF1*  Normalization: TMM (NOISeq) | LKB1_sig SQ: Mean of scaled expression for 16 genes across squamous tumors  LKB1_sig NSQ: Mean of scaled expression for 16 genes across non-squamous tumors | Var LKB1: If LKB1_sig > 0.5, Var LKB1 = 1; Else, Var$LKB1 = 0 |
| NRF2 score (unpublished) | *ABCB6, AKR1C1, AKR1C2, AKR1C3, TXNRD1, GCLM, SRXN1, PGD, TRIM16L, TRIM16, G6PD, OSGIN1, TALDO1, NQO1, NR0B1, UGDH, GSR, CYP4F3, AKR1B10*  Normalization: TMM (NOISeq) | NRF2_sig SQ: Mean of scaled expression for 19 genes across squamous tumors  NRF2_sig NSQ: Mean of scaled expression for 19 genes across non-squamous tumor | VarNRF2: If NRF2_sig > 0.5, score = 1; Else, VarNRF2 = 0 |
| Neuroendocrine differentiation score (unpublished) | *AGXT2L1, COL25A1, SLC14A2, ODC1, ENO3, ASCL1, CALCA, CALCB, RET, ZMAT4, MTMR7, SLC38A8, CELF3, KLK11, KLK13, KLK12, KLK14, SEC11C, BAALC, ERO1LB, CTNND2, UGT3A1*  Normalization: TMM (NOISeq) | Neuro_sig SQ: Mean of scaled expression for 22 genes across squamous tumors  Neuro_sig NSQ: Mean of scaled expression for 22 genes across non-squamous tumors | VarNeuro: If Neuro_sig > 0.5, score = 1; Else, VarNeuro = 0 |

PD-L1, programmed death ligand 1; TMB, tumor mutational burden; TMM, trimmed mean of M.

#### FICAN-OSCAR model

The full description of the model is available [here](https://www.synapse.org/#!Synapse:syn24180900/wiki/608579).

This model was used for all three sub-challenges as a linear product of the data matrix X and beta coefficients b. Gene expression signature for our custom gene panel (CUSTOM_FOPANEL) was estimated inside the Docker container using gene set variation analysis (GSVA), with parameter mx.diff=TRUE. Lastly, demographics such as sex or important clinical variables such as TMB and PD-L1 immunohistochemistry (IHC)-based model coefficients were derived as a symbiosis from widely reported trends in literature (see Table below) and the training data signals that we were able to utilize.

The model predicted a continuous value by assigning these regularized coefficients to beta vector (b) and then producing linear product Xb prediction over all three sub-challenges given the data matrix X. The resulting vector of length N (number of patients) was given as the final prediction. The coefficients were negative due to PFS and OS sub-challenges expecting lower values to correspond to better outcome (smaller hazard ratio), while BOR sub-challenge had the opposite direction (lower values corresponding to poor outcome). The directionality did not matter for final predictions, as the predictions in all sub-challenges were flipped around the null hypothesis for each sub-challenge (C-index_null_ = 0.5 in sub-challenges 1 and 2 and ROC-AUC_null_ = 0.5 in sub-challenge 3).

The Optimal Subset Cardinality Regression (*oscar*) L0-quasinorm penalized linear regression approach that was utilized has since been released as an independent R statistical software package in [https://github.com/Syksy/oscar/releases/tag/v0.6.1](https://nam02.safelinks.protection.outlook.com/?url=https%3A%2F%2Fgithub.com%2FSyksy%2Foscar%2Freleases%2Ftag%2Fv0.6.1&data=05%7C01%7CDoreen.Valentine%40bms.com%7Cf0ef1ed62b7e40a2e00208da7a05f1ed%7C71e34cb83a564fd5a2594acadab6e4ac%7C0%7C0%7C637956463207724623%7CUnknown%7CTWFpbGZsb3d8eyJWIjoiMC4wLjAwMDAiLCJQIjoiV2luMzIiLCJBTiI6Ik1haWwiLCJXVCI6Mn0%3D%7C3000%7C%7C%7C&sdata=dmzZdF%2BXXxYh5UElwpF6U6lxyRrK85v4AXQzmRahDrI%3D&reserved=0).^21,22^

FICAN-OSCAR model equation:

Y=–0.693×CUSTOM_FOPANEL–0.357×isTMBhigh–0.105×isMale–0.198×isSquamous–0.05×isSquamous&Above5PDL1–0.223×isEversmoker–0.105×isECOG

| **Name** | **Coefficient description** | **Estimate (b_i_)** | **Part of Challenge baseline models** | **Literature sources for curation** |
| --- | --- | --- | --- | --- |
| CUSTOM_FOPANEL | Panel of expression for *CD274*, *PDCD1*, *TIGIT*, *CXCL9*, and *CXCR6*, with score estimated using GSVA | -0.693 | No | Numerous sources supporting each gene candidate. For more details on the resources used, see Tables 1 to 3 here: <https://www.synapse.org/#!Synapse:syn24180900/wiki/608579> |
| isTMBhigh | Binary indicator if patient's TMB exceeded 243 | -0.357 | Yes | Carbone et al., estimate from Cristescu et al.^23,24^ |
| isMale | Binary indicator if patient was male | -0.105 | No | Conforti et al., Borghaei et al., Brahmer et al., training data in Prat et al.^25-28^ |
| isSquamous | Binary indicator if tumor was squamous | -0.198 | No | Comparisons between previous CheckMate trials with squamous or non-squamous NSCLC |
| isSquamous_above5PDL1 | Binary indicator for squamous tumors coupled with PD-L1 IHC >= 5% | -0.051 | Partly (PD-L1 IHC) | Carbone et al.^23^ |
| isEversmoker | Binary indicator if tobacco use was FORMER or CURRENT | -0.223 | No | Brahmer et al., Borghaei et al., Adachi et al, others^26,28,29^ |
| isECOG0 | Binary indicator if patient's ECOG status was equal to 0 | -0.105 | No | Brahmer et al., Borghaei et al., multiple other sources^26,28^ |

ECOG, Eastern Cooperative Oncology Group; GMT, gene matrix transposed file format; GSVA, gene set variation analysis; IHC, immunohistochemistry; NSCLC, non-small cell lung cancer; PD-L1, programmed death ligand 1; TMB, tumor mutational burden.

#### I-MIRACLE model

The full description of the model is available [here](https://www.synapse.org/#!Synapse:syn23746812/wiki/608744).

A 20-gene signature called the immunologic constant of rejection (ICR) that reflects Th-1 signaling activation (*IFNG, TBX21, CD8B, CD8A, IL12B, STAT1,* and *IRF1*), CXCR3/CCR5 chemokine ligands (*CXCL9, CXCL10, and CCL5*), cytotoxic effector molecules (*GNLY, PRF1, GZMA, GZMB, and GZMH*), and compensatory immune regulators (*CD274/PD-L1, PDCD1, CTLA4, FOXP3 and IDO1*) was reported previously (see Figure below).

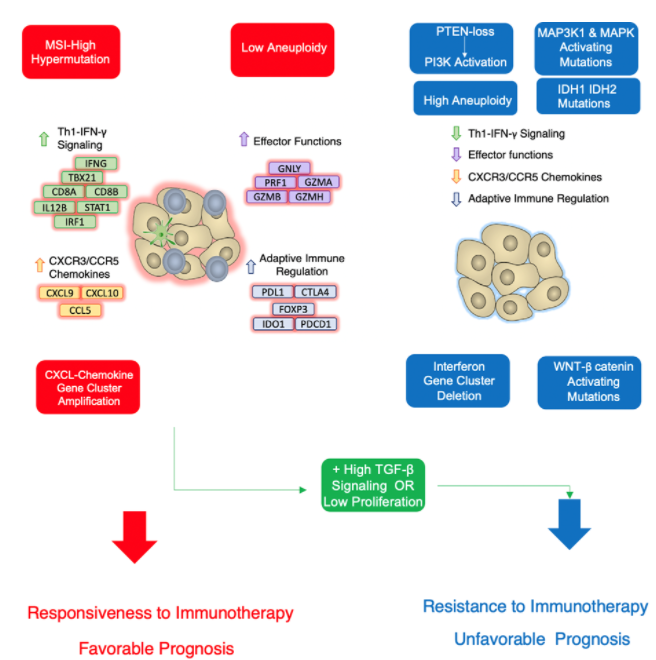

Adapted from Cancer Immunotherapy Principles and Practice, 2^nd^ Edition, p. 462), by D. Bedognetti et al., 2021, Place of Publication: Springer Publishing Company. Copyright 2021 by Springer Publishing Company. Adapted with permission.

IFN-γ, interferon gamma; MSI, microsatellite instability; TGF-β, transforming growth factor-beta.

The ICR is a 20-gene score that reflects the presence of a Th1/cytotoxic immune response, with a high ICR score predictive of response to immunotherapy and a low ICR score predictive of resistance to immunotherapy (see Figure below).^30,31^

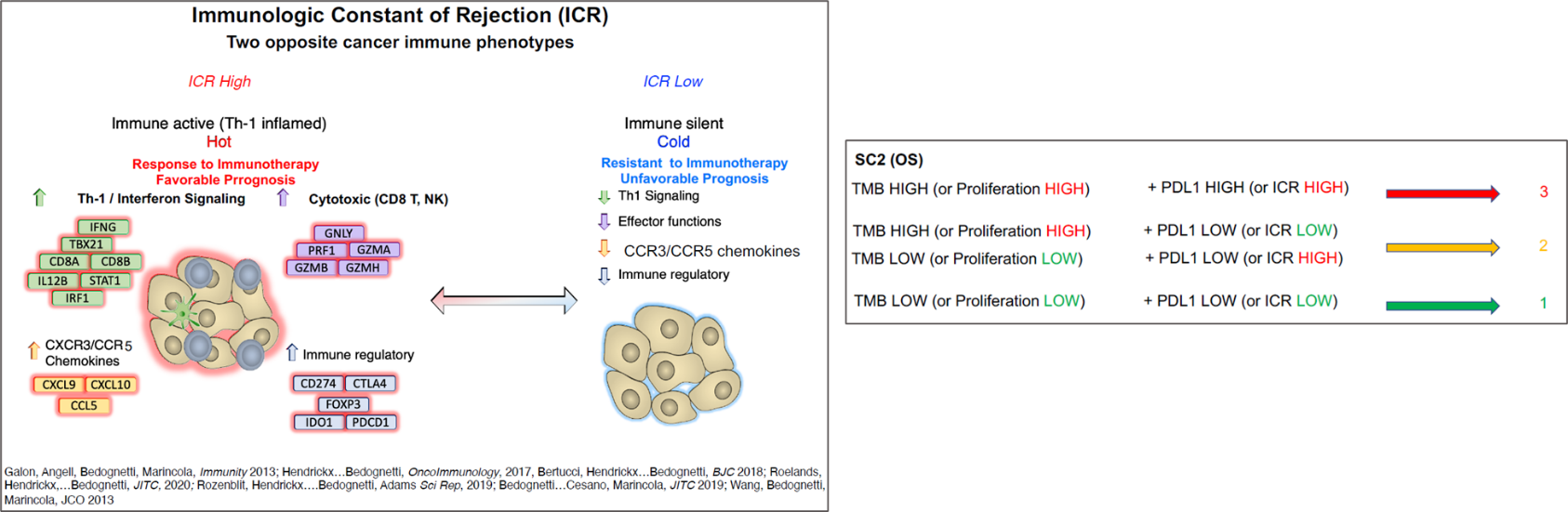
Adapted from Cancer Immunotherapy Principles and Practice, 2^nd^ Edition, p. 462), by D. Bedognetti et al., 2021, Place of Publication: Springer Publishing Company. Copyright 2021 by Springer Publishing Company. Adapted with permission.

The prediction model was built based on a categorical value (as scores) depending on the classification of TMB and PD-L1 as high and low as follows:

- **TMB**:
  - TMB values were classified as high and low based on the upper tertile as cut-off. A patient with TMB value higher or equal to the cut-off was considered TMB high, and below the cut-off was considered TMB low
  - When TMB was missing, the proliferation score was used instead,^32^ as it highly correlates with TMB in non-small cell lung cancer (NSCLC)
    - The gene expression profiles from LUAD (https://portal.gdc.cancer.gov/projects/TCGA-LUAD) and LUSC: (https://portal.gdc.cancer.gov/projects/TCGA-LUSC) cohorts from TCGA were used to perform a Pearson correlation with TMB. Five genes included in the proliferation signatures (*NUF2, NEK2, TPX2, KIF2C* and *MCM10*) were found to highly correlate with TMB (R = 0.48, p < 2.2e-16)
    - The enrichment score for this signature was calculated for all patients using the yaGST package (<https://github.com/miccec/yaGST>)
    - After obtaining the scores, tertiles were calculated to classify patients as TMB high or low depending on the value of upper tertile (66%)
    - Patients with missing TMB were classified as TMB high if their proliferation score was greater than or equal to the upper tertile and as TMB low otherwise
- **PD-L1**:
  - A patient with tumor cell PD-L1 expression ≥ 50% was considered PD-L1 high, and < 50 was considered PD-L1 low
  - When PD-L1 expression values were missing, ICR score was used instead
    - The enrichment score for this signature was calculated for all patients using the yaGST package
    - After obtaining the scores, tertiles were calculated to classify patients as PD-L1 expression high or low depending on the value of upper tertile (66%)
    - Patients with missing PD-L1 were classified as PD-L1 high if their ICR score was greater than or equal to the upper tertile and as PD-L1 low otherwise

For the OS and PFS sub-challenges, we built our model based on TMB, PD-L1, ICR, and proliferation signatures, as mentioned in the previous paragraphs. The output prediction file was submitted as a categorical value (score) as indicated in the table below:

| **Sub-challenge** | **TMB or proliferation status** | **PD-L1 or ICR status** | **I-MIRACLE score** |
| --- | --- | --- | --- |
| OS | TMB/proliferation high | PD-L1/ICR high | 3 |
|  | TMB/proliferation high | PD-L1/ICR low | 2 |
|  | TMB/proliferation low | PD-L1/ICR high |  |
|  | TMB/proliferation low | PD-L1/ICR low | 1 |
| PFS | TMB/proliferation high | PD-L1/ICR high | 3 |
|  | TMB/proliferation high | PD-L1/ICR low | 2 |
|  | TMB/proliferation low | PD-L1/ICR low | 1 |
|  | TMB/proliferation low | PD-L1/ICR high |  |

ICR, immunologic constant of rejection; OS, overall survival; PD-L1, programmed death ligand 1; PFS, progression-free survival; TMB, tumor mutational burden.

For the BOR sub-challenge, we built our model based on the calculation of ICR enrichment score. The output prediction file was submitted as a continuous ICR score.

The equation used to calculate the I-MIRACLE score was as follows:

$OS score = \delta\left( PDL1 \right)\times\delta\left( PDL1\geq50\% \right) + \left( 1-\delta\left( PDL1 \right) \right)\times\delta\left( ICR> \theta_{I} \right) + \delta\left( TMB \right)\times\delta\left( TMB> \theta_{T} \right)+\left( 1-\delta\left( TMB \right) \right)\times\delta\left( Proliferation > \theta_{P} \right) +1$

$PFS score =(\delta\left( TMB \right)\times\delta\left( TMB> \theta_{T} \right) +\left( 1-\delta\left( TMB \right) \right)\times\delta\left( Proliferation > \theta_{P} \right)) \times(2 +\delta\left( PDL1 \right)\times\delta\left( PDL1\geq50\% \right) + \left( 1-\delta\left( PDL1 \right) \right)\times\delta\left( ICR> \theta_{I} \right) +(\delta\left( TMB \right)\times\delta\left( TMB< \theta_{T} \right) +(1-\delta\left( TMB \right))\times\delta\left( Proliferation < \theta_{P} \right))$

$\theta_{I}:upper tertile of estimated ICR score from gene expression profiles of patients$

$\theta_{P}:upper tertile of estimated proliferation scores of patients$

$\theta_{T}:upper tertile of available TMB scores for patients$

$\delta\left( x \right)=1, if x is present or true$

$\delta\left( x \right)=0,if x is missing or false$

#### @jacob.pfeil model

The full description of the model is [here](https://www.synapse.org/#!Synapse:syn24305984/wiki/608243).

1. Description of the model

The @jacob.pfeil taux model used gene expression ratios to predict response to anti–PD-1 therapy in NSCLC. The motivation behind using gene expression ratios was to down-weight the effect of response markers by a factor proportional to resistance marker expression. This model only used gene expression data. TMB, histology, and clinical features were not used in developing the model. Previous methods for predicting response to checkpoint blockade typically focused on markers of response (ie, TMB, immune infiltrate, PD-L1 expression). The @jacob.pfeil taux approach has shown that superior predictive performance can be achieved if bulk gene expression markers of response are inversely weighted by paired resistance markers.

2. Model training

Two publicly available checkpoint response studies in NSCLC (GSE126044^33^ and GSE135222^34^) were identified. These studies were used because they were the only studies with gene expression and matched checkpoint blockade response data. Combining these studies resulted in 42 samples in total, where 30 samples were associated with progressive disease and 12 were associated with response. Gene ratios were then calculated and correlated with response. This identified ~300 gene ratios.

Investigation into gene functions found that genes in the numerator were associated with the adaptive immune response and genes in the denominator were associated with tumor intrinsic resistance mechanisms. While these features were obtained from the data, it appears that this approach identified expected markers of response while simultaneously accounting for tumor intrinsic resistance mechanisms.

These 300 gene ratios were then used to train a support vector machine (SVM) classifier. Given the limited training data, the @jacob.pfeil taux method used extensive bootstrapping and cross validation to identify parameters that were robust across thousands of resampled training and test data sets. During training, it was found that the non-linear radial basis function (RBF) kernel performed better than the linear kernel, suggesting that the decision boundary is not linearly separable until projected into higher dimensions by the RBF kernel.

3. Model versions

Two versions were submitted. V1 was submitted for PFS and OS sub-challenges. The BOR sub-challenge appeared to have swapped the labels for the prediction, so C3V1 was adjusted for this sub-challenge.

4. Steps in prediction

- Scale transcript per million (TPM) values using min-max scaler
- Determine class membership using previously trained SVM decision boundary
- Output binary prediction labels

5. Dockerizing the model

- Start by cloning this repository
- Build the Docker image that will contain the model with the following command:

docker build -t abbvie_taux:v1 .

6. Run the model

- Run the dockerized model

docker run \

-v $(pwd)/CM_026_formatted_synthetic_data_subset/:/data:ro \

-v $(pwd)/output:/output:rw \

abbvie_taux:v1

- The predictions generated are saved to /output/predictions.csv.

$ cat output/predictions.csv

patientID,prediction

p267,1

p315,0

p15,0

...

#### Netphar model

The full description of the model is available [here](https://www.synapse.org/#!Synapse:syn24241625/wiki/608681).

Previous studies suggested that both TMB and PD-L1 expression accounted for anti–PD-(L)1 response. The predictive power of PD-L1, however, is limited, particularly when a continuous expression level (higher than 1%) rather than a 50% cut-off is used.^28,35,36^ We considered two possible hypotheses:

- Hypothesis 1: PD-L1 contributes independently to the anti–PD-1 response, yet when compared with TMB, its contribution is relatively small. In this situation, a linear additive model can be fitted:

Y = b1 TMB + b2 PD-L1 (+ b3 sex+ b4 smoking + b5 gene_exp)

- Hypothesis 2: PD-L1 does not contribute independently to the anti–PD-1 response, but instead exerts its biological effect through an interplay with TMB. In this scenario, an interaction term should be considered:

Y = b1 TMB + b2 PD-L1 × TMB (+ b3 sex + b4 smoking + b5 gene_exp)

Given the extensive supporting evidence of TMB, we reasoned that b1 should always be larger than the other coefficients. To avoid overfitting when using a continuous variable, we binarized TMB by a threshold of ≥ 243. All the other continuous variables were scaled to have zero mean and unit variance. Values for the beta coefficients were estimated from the summary statistics of the CheckMate 026 publication.^23^ We found that our linear additive model achieved an accuracy slightly higher than using TMB alone when predicting BOR, suggesting a general validity of using linear models. However, the model performance was improved only when the coefficient of TMB was larger than that for PD-L1.

Due to the limited improvement using the linear additive model, we next tried the model with interaction terms:

Y = 10 × TMB_binarized_ + TMB_binarized_ × PD-L1

This model belongs to Hypothesis 2, which suggests that the immune suppression will be relieved (judged by PD-L1) only when the tumor can be recognized by immune cells as non-self (judged by TMB). The model outperformed all the baseline models in the three sub-challenges, with a primary score of 0.1708 in PFS prediction, 0.0666 in OS prediction, and 0.0175 in BOR prediction, respectively. This model can also be viewed as a decision tree, stating that TMB is necessary but not sufficient for triggering the response to immune checkpoint inhibitor (ICI), and PD-L1 becomes relevant only when TMB is high. On the other hand, the model is conservative on the TMB low branch, as all the predictions are zero. We also tried to include other covariates such as smoking, sex, immune genes, inflammatory signature, TCR evenness, and LKB1 expression, to further predict the TMB low branch. However, we did not see any further improvement when we optimized the model based on the leaderboard feedback (judged by our submissions in the leaderboard), suggesting that the anti–PD-1 response in TMB low patients is difficult to predict, most likely due to the convergent pathways that are important for both ICIs and chemotherapies in TMB low patients. Meanwhile, we also attempted to set PD-L1, instead of TMB, as the root of the tree model; however, it did not predict well, suggesting that PD-L1 is not the first cause.

#### Team TIDE model

The full description of the model is available [here](https://www.synapse.org/#!Synapse:syn25304416/wiki/609117).

1. Data preprocessing

- Immunedeconvolution (Immunedeconv) was performed on TPM without normalization
- TPM matrix was normalized using standard normalization before calculating the average expression of gene signatures
- TPM was log-transformed by $log2(x+1)$ and was normalized by quantile normalization. We then subtracted the average of the sample before calculating dysfunction and exclusion score using tumor immune dysfunction and exclusion (TIDE)
- Missing TMB was filled with the median of non-missing TMB

2. Feature selection

We used treatment-naive ICI clinical trial data from the TIDE database and late-stage chemotherapy patients of LUAD (https://portal.gdc.cancer.gov/projects/TCGA-LUAD), LUSC (https://portal.gdc.cancer.gov/projects/TCGA-LUSC), and skin cutaneous melanoma (SKCM) (https://portal.gdc.cancer.gov/projects/TCGA-SKCM) from TCGA as our training data.

We first calculated the C-index of survival of each feature within individual cohort and rank features according to the scoring metric:

$$primary metric= Mean (scale ({BM}_{nivo}^{SC})^{2})-Mean (scale ({BM}_{Chemo}^{SC})^{2})$$

Top features were selected in the model prediction, and the sources of these features are indicated in the table below.

| **Feature used in the model** | **Source** |
| --- | --- |
| TMB | From provided clinical data |
| PD-L1 | From provided clinical data |
| CTL | Calculated from TIDE (Supplementary Table 2) |
| SMOKE | From provided clinical data |
| Dysfunction | Calculated from TIDE (Supplementary Table 2) |
| Exclusion | Calculated from TIDE (Supplementary Table 2) |
| T.cell.CD4.non.regulatory_QUANTISEQ | Calculated from Immunedeconv |
| B.cell.naive_XCELL | Calculated from Immunedeconv |
| IFNG.Signature | Calculated from average expression of *IFNG, STAT1, IDO1, CXCL10, CXCL9*, and *HLA-DRA* (Supplementary Tables 2 and 3) |
| Antigen presentation by MHC-I | Calculated from average expression of *B2M,  HLA-A, HLA-B, HLA-C, HLA-E, HLA-F, HLA-G, HLA-H* and *TAP2* (list of genes curated from Wang S et al.^37^) |

CTL, cytotoxic T lymphocytes; Immunedeconv, immunedeconvolution; INFG, interferon gamma; MHC, major histocompatibility complex; PD-L1; programmed death ligand 1; TMB, tumor mutational burden.

3. Methods for PFS sub-challenge

According to the prior knowledge of this trial from the Challenge committee, TMB and PD-L1 are baseline markers for PFS. We calculated the rank product using TMB and PD-L1 as a predictor for this sub-challenge. The final prediction was calculated using the below function:

$$prediction=rank({TMB)}^{2}\times rank (PDL1)$$

Build docker image

docker build -f Dockerfile_PFS -t awesome-antipd1-q1-model:vv

4. Methods for OS sub-challenge

Seven immune features (TMB, PD-L1, CTL, T.cell.CD4.non.regulatory_QUANTISEQ, B.cell.naive_XCELL, Exclusion, and IFNG.Signature) and smoking status were integrated in the OS prediction. Because the concordance index was measured between the rank of predicted OS and observed OS, we first calculated the aggregated rank of the seven immune features using the Robust Rank Aggregation method,^38^ and then adjusted the aggregated ranking according to the patient’s smoking status. We assumed the order of TMB, PDL1, CTL, T.cell.CD4.non.regulatory_QUANTISEQ, and B.cell.naive_XCELL were in the same direction with OS, while the exclusion score was in the reverse direction. After calculating the aggregated ranking of those seven immune features, we then increased the rank by 2 for current smokers and by 1 for former smokers. We also decreased the rank by 1 for non-smokers. Assuming smoking was a key factor, the ranking adjustment could further differentiate the OS. The final ranking was our prediction of OS.

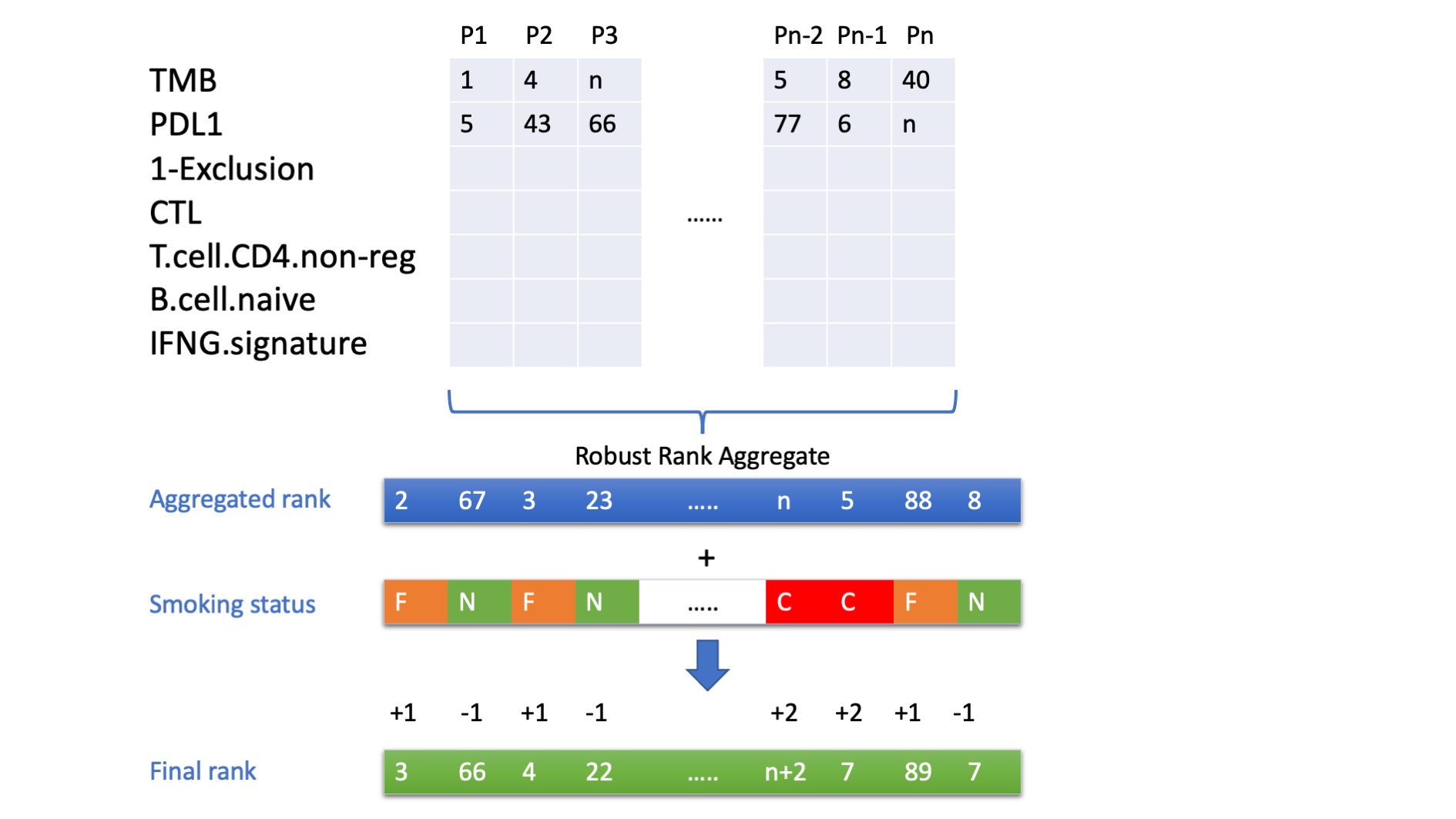

Smoking status F: former smoker; N: never a smoker; C; current smoker.

5. Methods for BOR sub-challenge

PD-L1 expression and TMB are informative of the tumor immune environment that determines the anti-tumor effect of ICI treatments. Hence, we developed four different sub-models depending on their PD-L1 and TMB levels. They were PD-L1^high^TMB^high^, PD-L1^low^TMB^high^,
PD-L1^high^TMB^low^, and PD-L1^low^TMB^low^.

Tumors with PD-L1^low^TMB^high^ status indicate the enrichment of cytotoxic T cells, which associates with the response to ICI.^39,40^ However, if these T cells are in a dysfunctional state, tumors will be resistant to the immunotherapy treatment.^39^ Thus, we applied the TIDE model^41^ to quantify the T-cell dysfunctional level of these tumors and predict their response to ICI by the negative dysfunction score, which was denoted by -Dys(x).

We hypothesized that tumors with high PD-L1 but low TMB (PD-L1^high^TMB^low^) evade immune cell killing by either excluding T-cell infiltration or having antigen presentation defect. Therefore, we considered both the T-cell exclusion score and antigen presentation level when estimating the response to ICI.^42^ Briefly, we calculated the T-cell exclusion score by the TIDE model^41^ (Exc(x)) and evaluated the antigen presentation level (Ap(x)). We then predicted the response to ICI by the average value of these two scores.

Next, considering that tumors with PD-L1^high^TMB^low^ status tend to have more dysfunctional T cells, we also predicted the response of these tumors by the T-cell dysfunction score, but in the opposite direction to tumors with PD-L1^low^TMB^high^ status.

Lastly, tumors with PD-L1^low^TMB^low^ status might be the most resistant subgroup to the ICI. In this subgroup, we assumed that only tumors with relatively high PD-L1 levels and antigen presentation incidence can trigger response to the ICI. According to this hypothesis, we predicted tumors with PD-L1^low^TMB^low^ status by the average value of PD-L1 level and antigen presentation level. Taking those together, we built a conditional model with the following formula to predict response to the ICI.

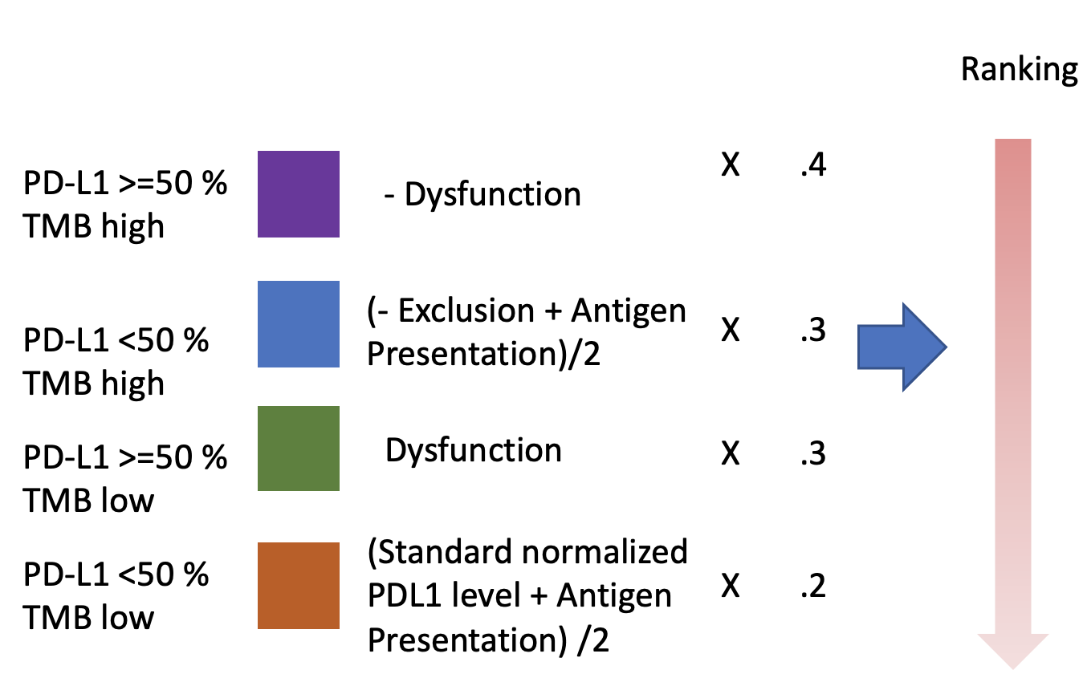

Build docker image

docker build -f Dockerfile_response -t awesome-antipd1-q3-model:vv

### Supplementary Methods 2. Model submission by participants and model evaluation

DREAM Challenge participants had a quota of five valid submissions during the validation phase for each sub-challenge question. The most recent valid submission (ie, the final model) was used to determine the top-performing model for each sub-challenge. Participants received an estimate of their score computed by taking the mean of 10 bootstrapped samples. Estimates were provided for both the primary metric and tie-breaking metrics, as well as performance in the chemotherapy arm. The custom metrics used to evaluate model performance were designed to identify models that differentiated patient benefit in the nivolumab arm but did not differentiate patient benefit in the chemotherapy arm.

**Open phase
4 weeks**

**Synthetic validation phase
8 weeks**

**Validation phase**

**Final round**

**Scoring period**

Nov 17^th^, 2020

Dec 15^th^, 2020
Synthetic data queues open

Feb 10^th^, 2021
Validation data queues open

Feb 25^th^, 2021
Validation data queues close

Mar 23^rd^, 2021
Top performers announced

Dec 4^th^, 2020 Webinar

The distributions of participants’ primary metric scores were computed via bootstrapping. For each sub-challenge question, a Bayes factor (K) was computed for each team’s model using the top-performing model as the reference. The Bayes factor was used to determine a group of statistically tied teams (eg, K < 3), and the tie-breaking metric was then applied to this group. The tie-breaking metric for each sub-challenge was the basal metric applied in the nivolumab arm of the CheckMate 026 trial.

### Supplementary Methods 3. Pathway Analysis of Gene sets

All the top-performing teams who generated models that relied on gene expression data were asked to provide a ranking for each gene (total of 20,000 genes each) according to the importance of the gene in the model. The ranks from all teams were collated and the union of the top 50 genes from each of the models was selected for downstream analysis. The expression levels of these genes in the RNAseq data included in CheckMate 026 dataset were noted. All gene expression data were z-score normalized. The selected gene set was further expanded to include any additional genes whose expression levels had high correlation (correlation > 0.85) with the initially selected genes. Unsupervised hierarchical clustering of the gene expression data of the selected gene set was performed and the resulting gene clusters were visualized using the pheatmap R package (the dendrogram was cut at a height of 23.4 to select the gene clusters). A hypergeometric test using the gProfiler2 R package was carried out to explore the enrichment of pathways from [Gene Ontology](http://geneontology.org/) (GO), [Kyoto Encyclopedia of Genes and Genomes](https://www.genome.jp/kegg/) (KEGG), [Reactome](https://reactome.org/) (REAC), [WikiPathways](https://www.wikipathways.org/index.php/WikiPathways) (WP), [TRANSFAC](https://genexplain.com/transfac/) (TF), [MiRTarBase](https://mirtarbase.cuhk.edu.cn/~miRTarBase/miRTarBase_2019/php/index.php) (MIRNA), [Human Protein Atlas](https://www.proteinatlas.org/) (HPA), [CORUM](http://mips.helmholtz-muenchen.de/corum/) (database of mammalian protein complexes), and [Human Phenotype Ontology](https://hpo.jax.org/app/) (HP) in the selected gene set. Pathway terms with a term size less than 100 from the KEGG and REAC databases enriched in the selected gene set were explored further.

### Supplementary Table 1. Candidate predictors available to participating teams

| **ID** | **Label** | **Class** | **Levels** | **Note** |
| --- | --- | --- | --- | --- |
| patientID | unique deidentified patient ID | character |  |  |
| SEX | Sex | factor | F; M |  |
| AAGE | Derived Age at Consent | numeric |  |  |
| CRFHIST | CRFHIST: CRF HISTOLOGY | factor | NON-SQUAMOUS; SQUAMOUS | Important variable histology |
| TOBACUSE | Tobacco Use | factor | CURRENT; FORMER; NEVER; UNKNOWN | Important variable |
| ECOGPS | ECOGPS: PERFORMANCE STATUS (ECOG) | numeric |  | Important variable ECOG |
| PDL1 | PDL1 IHC Percent | numeric |  | Important variable |
| TMB | Tumor Mutation Burden | numeric |  | Important variable |
| TCR_Shannon | TCR Shannon's Entropy | numeric |  |  |
| TCR_Richness | TCR Richness | integer |  |  |
| TCR_Evenness | TCR Evenness | numeric |  |  |
| BCR_Shannon | BCR Shannon's Entropy | numeric |  |  |
| BCR_Richness | BCR Richness | integer |  |  |
| BCR_Evenness | BCR Evenness | numeric |  |  |

BCR, B-cell receptor; CRF, case report form; ECOG, Eastern Cooperative Oncology Group; F, female; ID, identifier; IHC, immunohistochemistry; M, male; PDL1, programmed death ligand 1; TCR, T-cell receptor; TMB, tumor mutational burden.

### Supplementary Table 2. Comparator models: Published and baseline models used for benchmarking

Eleven published gene expression-based predictors of patients’ responses to ICI immunotherapy are described below along with two baseline models using clinical traits. We made use of the functions available at *easier* R package to compute scores for all predictors from bulk RNA-seq with TPM values.^1^ Predictors’ larger values were associated with higher likelihood of response to immunotherapy.

| **Model used for benchmarking** | **Short model description** |
| --- | --- |
| *Published models* | |
| ICB_tcellinflamed^10^ | This signature was defined by an 18-gene set derived from a penalized logistic regression to predict BOR to anti–PD-1 therapy. To compute the score, the expression of each gene was first normalized using an 11-gene housekeeping set showing low variance across many tumor samples from multiple cancer types. Using the regression coefficients, the final score was computed as a weighted sum of the normalized expression values (log2[TPM + 1]). |
| ICB_msi^17^ | First, an initial set of 4769 genes was found differentially expressed between microsatellite instability (MSI) and microsatellite stability (MSS) samples. From all the MSI-related gene pairs formed by these genes, 1,654,739 gene pairs were defined based on their high frequency in the MSI samples. To reduce the number of gene pairs, only the gene pairs with high frequency difference values (ie, more frequent in either the MSI or MSS class) were considered. The MSI status signature was then defined based on a set of 10 MSI-related gene pairs that best predict MSI status in right-sided colon cancer. For each sample, a score was assigned to each gene pair (gene $i$, gene $j$) based on the following function: f_ij_ = 1 if ${exp}_{i}>{exp}_{j}$ and f_ij_ = 0 if ${exp}_{i}<{exp}_{j}$where ${exp}_{i}$ and ${exp}_{j}$correspond to the expression (log2[TPM + 1]) of genes $i$ and $j$. The final score for each sample was computed as the sum of the result of the logical relations between the gene pairs. When there were missing pairs, the final score was normalized to the number of available gene pairs. |
| ICB_ifnγ^10^ | This signature was based on IFNγ-signaling-related genes. First, the 10 top-ranked genes were selected as the one separating responders versus non-responders to anti–PD-1 therapy in a melanoma cohort. Then, among these 10 genes, the genes significantly associated with either positive response or progression-free survival in an independent dataset were selected. The resulting six genes were considered the signature genes, and the score for each sample is computed as the average of their expression (log2[TPM + 1]). |
| ICB_cyt^6^ | This signature was quantified using the geometric mean of the gene expression (TPM + 0.01) of two cytolytic effectors, granzyme A and perforin. These two genes were found overexpressed in activated CD8+ T cells ^43^ and upon treatment with ICI ^44,45^. |
| ICB_chemokine^8^ | This signature was based on the PC1 score obtained by applying PCA to the normalized (z-score) expression (log2[TPM + 1]) of a 12-chemokine gene set associated with immunity and inflammation, which predicts the presence of lymph nodal structures in melanoma. In our computation of this score, we set the sign of the score according to a high positive correlation with the rest of the predictors (all except ICB_impres and ICB_msi). This was done because the sign of the first principal component is defined at random. |
| ICB_explS^10^ | First, a 28-gene set consisting of genes that showed high correlation with ICB_ifnγ signature genes in an initial melanoma dataset was generated. This signature included genes related to T-cell cytolytic activity, antigen presentation, and chemokine production. Then, the signature was compressed to an 18-gene set that showed significant association with either positive response or progression-free survival in an independent dataset. The final score for each sample was computed as the average of the gene expression of this 18 gene-set (log2[TPM + 1]). |
| ICB_davoliis^9^ | This signature was defined using the expression of seven molecular markers specific for cytotoxic CD8+ T cells and NK cells. To compute the score, first the expression of each gene (log2[TPM + 1]) was rank-normalized across samples, ie, the expression of each gene is replaced by its fractional rank (rank divided by the total number of genes). Then, the final score for each sample was calculated as the average of the fractional rank of all signature genes. |
| ICB_rohis^7^ | This signature was defined as the geometric mean of the expression (TPM + 0.01) of a set of 41 immune-related genes that were linked to the characterization of tissue destruction, hypothesized to be the common ground (so-called “immunologic constant of rejection”) shared by different immune processes, including tumor rejection ^46,47^. This gene signature included genes related to cytotoxic factors, HLA molecules, IFNγ pathway, specific chemokines, and adhesion molecules. |
| ICB_impres^48^ | First, a manually curated list of 28 immune checkpoint genes was used to generate 294 gene pairs, where at least one of the genes was associated with anti–CTLA-4 or anti–PD-1 immunotherapy. The IMPRES signature was then defined based on the subset of 15 gene pairs that best discriminate responders versus nonresponders in neuroblastoma. For each sample, a score was assigned to each gene pair (gene $i$, gene $j$) based on the following function: f_ij_ = 1 if ${exp}_{i}>{exp}_{j}$ and f_ij_ = 0 if ${exp}_{i}<{exp}_{j}$. Where ${exp}_{i}$ and ${exp}_{j}$correspond to the expression (log2[TPM + 1]) of genes $i$ and $j$. The final score for each sample was computed as the sum of the result of the logical relations between the 15 immune checkpoint gene pairs. When there were missing pairs, the final score was normalized to the number of available gene pairs. |
| ICB_rir^11^ | This model was defined by a set of three gene signatures describing major mechanisms of evasion and resistance to anticancer immunity: T-cell exclusion, resistance to ICI immunotherapy, and impaired T-cell–mediated killing.  The first one was the T-cell exclusion signature (“exc.down” in the original paper), which included genes expressed by malignant cells that are strongly positively correlated with T-cell infiltration, The second one was the resistance to ICI immunotherapy signature (“trt.down”) encompassing genes downregulated in malignant cells from tumors resistant to immunotherapy. The third one was related to impaired T-cell–mediated killing (“fnc.down”) encompassing genes that were downregulated in malignant cells which were defined resistant to T-cell killing.  Each signature score was calculated on the basis of an approach to hold a high signal-to-noise ratio by generating 1000 random permutations of the gene expression signature, so that each permutation preserved the overall signature distribution, to obtain 1000 random scores. The signature score was then defined as the mean-centered original gene expression (log2[TPM + 1]) minus the average of the random scores. The final score for each sample was calculated by adding the “fnc.down” score to the average of the scores for “exc.down” and “trt.down” signatures. |
| ICB_tls^12^ | This signature was based on a set of nine genes upregulated in melanoma tumors infiltrated by CD8+ T cells and CD20+ B cells that showed presence of tertiary lymphoid structures. The scores were computed using the geometric mean of its signature genes (TPM + 1). |
| ICB_TIDE^41^ | The TIDE web platform integrated omic data from 998 tumors from clinical studies with ICIs, as well as non-immunotherapy tumor profiles, and data from CRISPR screens. The platform takes gene set or expression profiles as input and prioritizes data for a user-input gene set, biomarker evaluation for a custom biomarker gene set, and biomarker consensus to predict responses to ICIs from gene expression profiles. |

| *Baseline models* | |
| --- | --- |
| TMB baseline^23^ | This model consisted of a Boolean variable indicating whether a sample TMB was in the top tertile within its cohort. |
| PD-L1 baseline^23^ | This model consisted of a Boolean variable indicating whether a sample PD-L1 expression was ≥ 50%. |

BOR, best overall response; CRISPR, clustered regularly interspaced short palindromic repeats; CTLA-4, cytotoxic T-lymphocyte-associated protein 4; HLA, human leukocytes antigen; ICB_cyt, immune cytolytic activity; ICB_davoliis, Davoli’s immune signature; ICB_impres, immune predictive score; ICB_msi, microsatellite instability status; ICB_rir, repressed immune resistance; ICB_rohis, Roh’s immune signature; ICB_tls, tertiary lymphoid structure; ICI, immune checkpoint inhibitor; IFN, interferon; NK, natural killer; PC, principal component; PCA, principal component analysis; PD-1, programmed death 1; PD-L1, programmed death ligand 1; TMB, tumor mutational burden; TPM, transcript per million.

### Supplementary Table 3. Components of the published gene signatures detailed in Supplementary Table 2

| **Published predictor (reference)** | **Gene signature** |
| --- | --- |
| Immune cytolytic activity  (ICB_cyt)^6^ | GZMA, PRF1 |
| Immunopredictive score (ICB_impres)^48^ | PDCD1, CD27, CTLA4, CD40, CD86, CD28, CD80, CD274, TNFRSF14, TNFSF4, TNFSF9, C10orf54, HAVCR2, CD200, CD276 |
| Roh immune score (ICB_rohis)^7^ | GZMA, GZMB, PRF1, GNLY, HLA-A, HLA-B, HLA-C, HLA-E, HLA-F, HLA-G, HLA-H, HLA-DMA, HLA-DMB, HLA-DOA, HLA-DOB, HLA-DPA1, HLA-DPB1, HLA-DQA1, HLA-DQA2, HLA-DQB1, HLA-DRA, HLA-DRB1, IFNG, IFNGR1, IFNGR2, IRF1, STAT1, PSMB9, CCR5, CCL3, CCL4, CCL5, CXCL9, CXCL10, CXCL11, ICAM1, ICAM2, ICAM3, ICAM4, ICAM5, VCAM1 |
| Chemokine signature  (ICB_chemokine)^8^ | CCL2, CCL3, CCL4, CCL5, CCL8, CCL18, CCL19, CCL21, CXCL9, CXCL10, CXCL11, CXCL13 |
| Davoli immune signature (ICB_davoliis)^9^ | CD247, CD2, CD3E, GZMH, NKG7, PRF1, GZMK |
| IFNγ signature (ICB_ifny)^10^ | IFNG, STAT1, CXCL9, CXCL10, IDO1, HLA-DRA |
| Immune expanded signature (ICB_expIS)^10^ | GZMB, GZMK, CXCR6, CXCL10, CXCL13, CCL5, STAT1, CD3D, CD3E, CD2, IL2RG, NKG7, HLA-E, CIITA, HLA-DRA, LAG3, IDO1, TAGAP |
| T-cell–inflamed microenvironment signature (ICB_tcellinflamed)^10^ | Signature: CCL5, CD27, CD274 (PD-L1), CD276 (B7-H3), CD8A, CMKLR1, CXCL9, CXCR6, HLA-DQA1, HLA-DRB1, HLA-E, IDO1, LAG3, NKG7, PDCD1LG2 (PDL2), PSMB10, STAT1, TIGIT  Housekeeping:  STK11IP, ZBTB34, TBC1D10B, OAZ1, POLR2A, G6PD, ABCF1, NRDE2, UBB, TBP, SDHA |

| Microsatellite instability status  (ICB_msi)^17^ | HNRNPL, MTA2, CALR, RASL11A, LYG1, STRN3, HPSE, PRPF39, CCRN4L, AMFR, CDC16, VGF, SEC22B, CAB39L, DHRS12, TMEM192, BCAS3, ATF6, GRM8, DUSP18 |
| --- | --- |
| Tertiary lymphoid structures signature (ICB_tls)^12^ | CD79B, CD1D, CCR6, LAT, SKAP1, CETP, EIF1AY, RBP5, PTGDS |
| Repressed immune resistance (ICB_rir)^11^ | The list of genes is made available by original publication, and it can be found by following the instructions here: <https://github.com/livnatje/ImmuneResistance/tree/master/Data> |

### Supplementary Table 4. Characteristics at baseline of all the patients who underwent randomization in CheckMate 026^23^

| **Characteristics** | **Nivolumab  (N = 271)** | **Chemotherapy (N = 270)** |
| --- | --- | --- |
| Age, years  Median (range) | 63 (32–89) | 65 (29–87) |
| Age ≥ 75 years, n (%) | 30 (11) | 32 (12) |
| Female sex, n (%) | 87 (32) | 122 (45) |
| Disease stage, n (%)  Stage IV  Recurrent  Not reported | 255 (94)  16 (6)  0 | 244 (90)  25 (9)  1 (< 1) |
| ECOG PS,^a^ n (%)  0  1  ≥ 2  Not reported | 85 (31)  183 (68)  2 (1)  1 (< 1) | 93 (34)  174 (64)  3 (1)  0 |
| Smoking status, n (%)  Never smoked  Former smoker  Current smoker  Unknown | 30 (11)  186 (69)  52 (19)  3 (1) | 29 (11)  182 (67)  55 (20)  4 (1) |
| Previous systemic therapy, n (%)  Adjuvant  Neoadjuvant | 22 (8)  5 (2) | 25 (9)  4 (1) |
| Previous radiotherapy, n (%) | 102 (38) | 107 (40) |
| Tumor histologic findings, n (%)  Squamous-cell carcinoma  Nonsquamous cell carcinoma | 66 (24)  205 (76) | 64 (24)  206 (76) |
| Selected site of metastatic lesions, n (%)  Brain  Liver | 33 (12)  54 (20) | 36 (13)  36 (13) |
| Sum of target-lesion diameters, mm  Median (range) | 82 (14–218) | 68 (15–272) |
| PD-L1 expression level, n (%)  ≥ 5%  ≥ 50% | 208 (77)  88 (32) | 210 (78)  126 (47) |
| Tumor mutational burden,^b^ n (%)  Low  Medium  High | 62 (39.2)  49 (31.0)  47 (29.7) | 41 (26.6)  53 (34.4)  60 (39.0) |

ECOG PS, Eastern Cooperative Oncology Group performance status; PD-L1, programmed death ligand 1.

^a^The ECOG PS score is assessed on a 5-point scale, with higher numbers indicating greater disability. Patients were required to have an ECOG PS score of 0 or 1 during screening. However, at baseline, the score had worsened to 2 in five patients and was not reported in one patient. ^b^Tumor mutational burden was available in 158 patients in the nivolumab arm and 154 patients in the chemotherapy arm.

### Supplementary Table 5. Characteristics at baseline of all the patients who underwent randomization in CheckMate 227^16^

| **Characteristics** | **Nivolumab + ipilimumab  (N = 583)** | **Chemotherapy (N = 583)** |
| --- | --- | --- |
| Median age, years | 64 | 64 |
| Female, % | 33 | 34 |
| ECOG PS, %  0  1  ≥ 2  Not reported | 35  65  < 1  0 | 33  66  1  < 1 |
| Smoking status, %  Never smoked  Current/former smoker  Unknown | 14  85  1 | 13  86  1 |
| Histology, %  Squamous  Nonsquamous | 28  72 | 28  72 |
| PD-L1 expression, %  < 1%  ≥ 1% | 32  68 | 32  68 |

ECOG PS, Eastern Cooperative Oncology Group performance status; PD-L1, programmed death ligand 1.

### Supplementary Table 6. Model performance across sub-challenges

|  | **PFS sub-challenge** | | **OS sub-challenge** | | **BOR sub-challenge** | |
| --- | --- | --- | --- | --- | --- | --- |
| **Model** | **Primary metric** | **Bayes factor** | **Primary metric** | **Bayes factor** | **Primary metric** | **Bayes factor** |
| Aginome_Amoy | 0.0185 | 142 | 0.0048 | 11 | 0.0524 | 1 |
| cSysImmunoOnco | 0.0354 | 44 | -0.0166 | 28 | 0.0545 | 0 |
| DukeLKB1 | 0.0506 | 24 | 0.0316 | 2 | -0.0838 | 6 |
| FICAN-OSCAR | 0.0257 | 62 | 0.0456 | 1 | 0.0389 | 1 |
| I-MIRACLE | 0.0867 | 24 | 0.0496 | 0 | 0.0047 | 3 |
| @jacob.pfeil | -0.0022 | 10 | 0.0721 | 1 | -0.0055 | 3 |
| Netphar | 0.1900 | 0 | 0.0009 | 7 | -0.0521 | 6 |
| Team TIDE | 0.0177 | 82 | 0.0164 | 3 | 0.0492 | 1 |
| ICB_msi | 0.0212 | 18 | 0.0101 | 3 | -0.0033 | 3 |
| ICB_chemokine | -0.0003 | 82 | 0.0041 | 6 | -0.0347 | 4 |
| ICB_cyt | -0.0020 | 110 | 0.0060 | 5 | 0.0007 | 3 |
| ICB_davoliis | -0.0115 | 142 | 0.0032 | 5 | -0.0301 | 5 |
| ICB_expIS | -0.0133 | 124 | 0.0036 | 5 | -0.0787 | 8 |
| ICB_ifny | 0.0049 | 82 | 0.0091 | 4 | -0.0353 | 5 |
| ICB_impres | 0.0004 | 199 | -0.0074 | 9 | -0.0042 | 3 |
| ICB_rir | -0.0223 | 199 | -0.0162 | 16 | -0.0896 | 11 |
| ICB_rohis | -0.0236 | 249 | -0.0056 | 7 | -0.1147 | 10 |
| ICB_tcellinflamed | -0.0117 | 82 | 0.0179 | 3 | -0.0666 | 7 |
| ICB_TIDE | 0.0024 | 66 | 0.0132 | 3 | -0.0522 | 5 |
| ICB_tls | -0.0355 | 499 | -0.0185 | 17 | -0.2104 | 21 |
| PD-L1 BL | 0.0018 | 124 | -0.0006 | 8 | -0.1019 | 8 |
| TMB BL | 0.0700 | 42 | 0.0067 | 4 | -0.0188 | 3 |

BL, baseline; BOR, best overall response; OS, overall survival; PD-L1, programmed death ligand 1; PFS, progression-free survival; TMB, tumor mutational burden.

### Supplementary Figure 1. Computing of the primary metric for each sub-challenge

1. **Computation of basal metric for each
   sub-challenge**

- The basal metrics are transformed to adjust their range and midpoint from (0,1, mid=0.5) to (-1, 1, mid=0) by the function $scaled (BM)=2\times(BM-0.5)$
- The scaled basal metric is then squared
- The primary metric is then calculated as the DSS basal metrics between the nivolumab and chemotherapy arms
- *PFS sub-challenge:* PFS Harrel’s concordance index with values termed ${BM}_{NIVO}^{PFS}$ and ${BM}_{CHEMO}^{PFS}$
- *OS sub-challenge:* OS Harrel’s concordance index with values termed ${BM}_{NIVO}^{OS}$ and ${BM}_{CHEMO}^{OS}$
- *BOR sub-challenge:* ROC-AUC with values termed ${BM}_{NIVO}^{BOR}$ and ${BM}_{CHEMO}^{BOR}$

2. **Basal metric transformation**
3. **Primary metric calculation**

AUC, area under the curve; BM, basal metric; BOR, best overall response; DSS basal metrics, difference in squared scaled basal metrics; OS, overall survival; PFS, progression-free survival; ROC, receiver operating characteristic.

### Supplementary Figure 2. Model performance

**(A) Submitted models’ performance in CheckMate 026 (data from the chemotherapy arm and nivolumab arm) and CheckMate 227 (data from the chemotherapy arm and nivolumab + ipilimumab arm). (B) Alternative model performance in CheckMate 026 and CheckMate 227. We computed performance using a more traditional measure whereby a three-term Cox proportional-hazards model was fitted with treatment arm, team predictions, and the arm-by-prediction interaction terms. A model performance can be measured by the estimated coefficient of the interaction terms, with values near zero showing that the model performed randomly. This coefficient is the ratio of the hazard ratios in the model. The plot shows the log_2_ of the coefficient and the log_2_ of the 95% confidence interval of this coefficient estimate.**

AUC, area under the curve; BOR, best overall response; DSS, difference in squared scaled basal metric; HR, hazard ratio; ICI, immune checkpoint inhibitor; OS, overall survival; PFS, progression-free survival; TMB, tumor mutational burden.

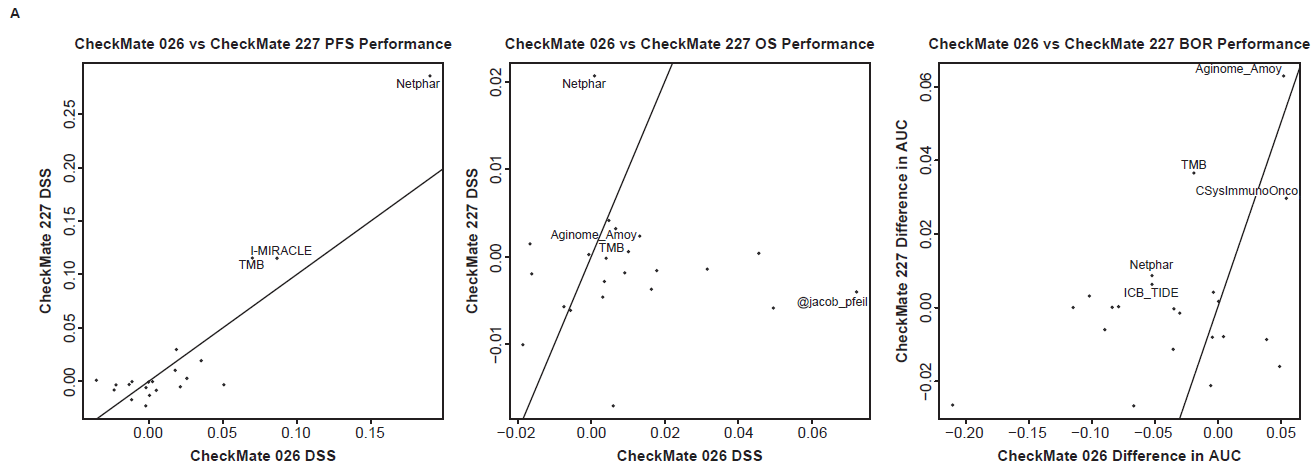

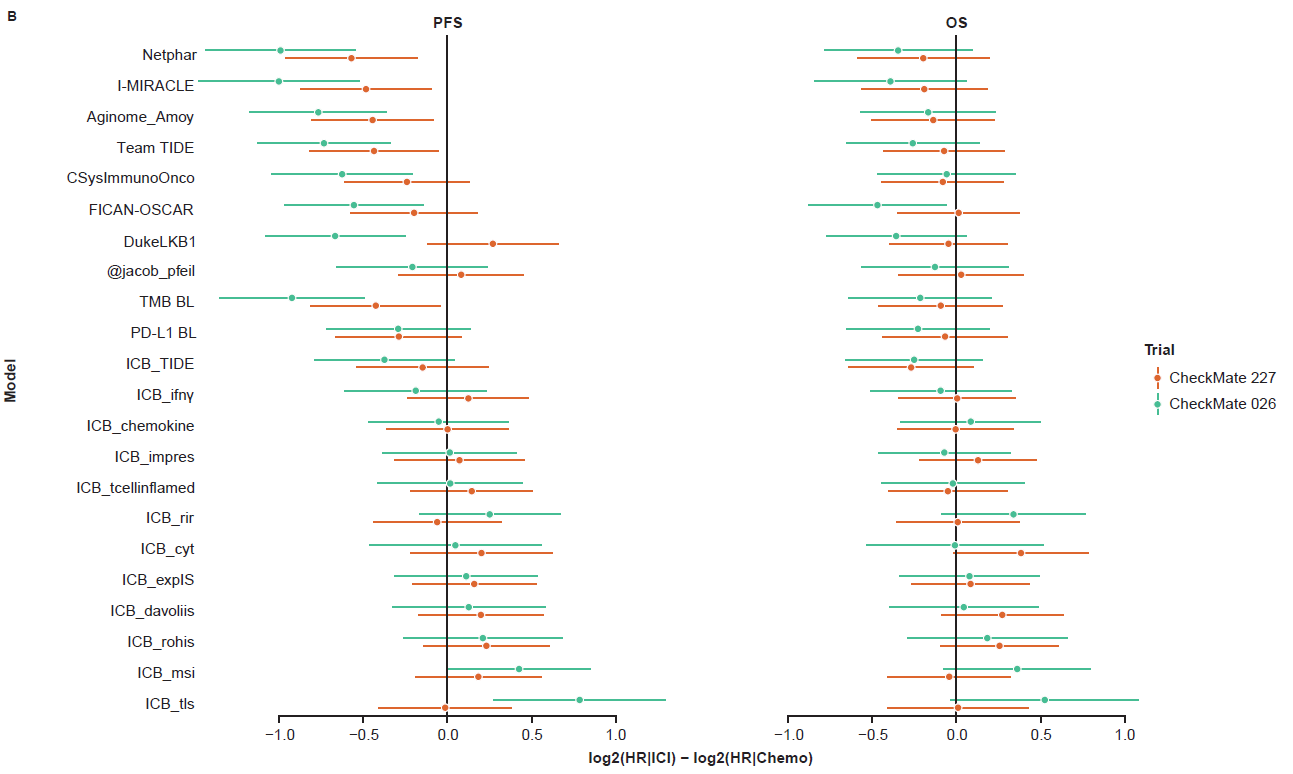

### Supplementary Figure 3. Gene signatures

**(A) Heatmap showing top genes ranked by models that relied on gene expression data. Any genes that did not receive a rank from any of the models were assigned a score of 30 and colored blue in the heatmap. (B) Heatmap showing correlation of gene expression among the selected gene sets and the clusters formed by hierarchical clustering of the gene expression data. (Ci-iii) Manhattan plots showing enrichment of pathways from various databases in Clusters 1, 2, and 3, respectively. (Di-iii) Tables showing the top pathways from KEGG or REAC for each of the identified clusters.**

BOR, best overall response; BP, biological processes; CC, cellular component; CORUM, database of mammalian protein complexes; CR, complete response; HP, Human Phenotype Ontology; HPA, Human Protein Atlas; KEGG, Kyoto Encyclopedia of Genes and Genomes; LOOCV, leave one out cross-validation; MF, molecular function; MIRNA, MiRTarBase; NE, not estimable; PD, progressive disease; PR, partial response; REAC, REACTOME; SD, stable disease; TF, TRANSFAC; WP, WikiPathways.

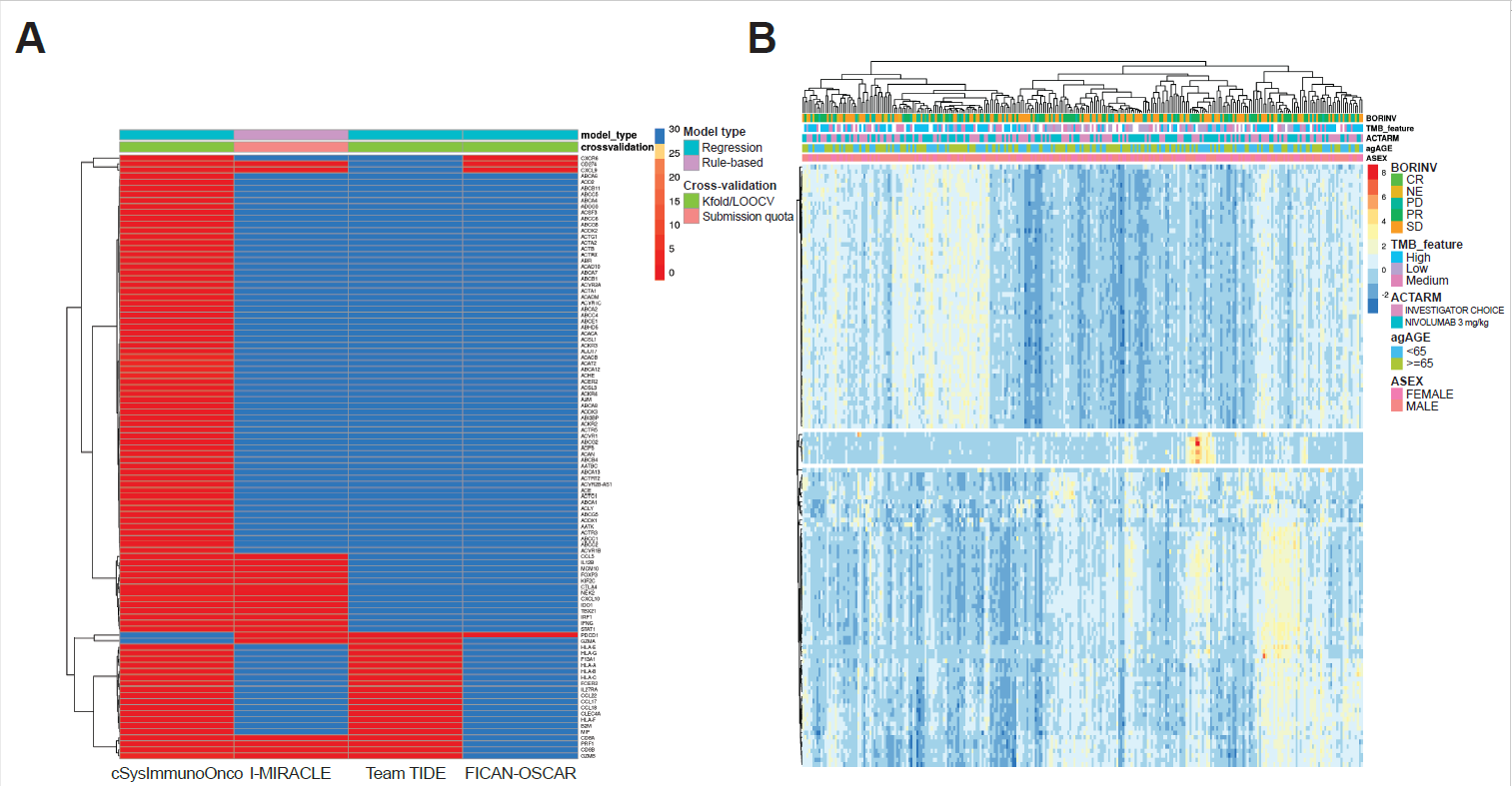

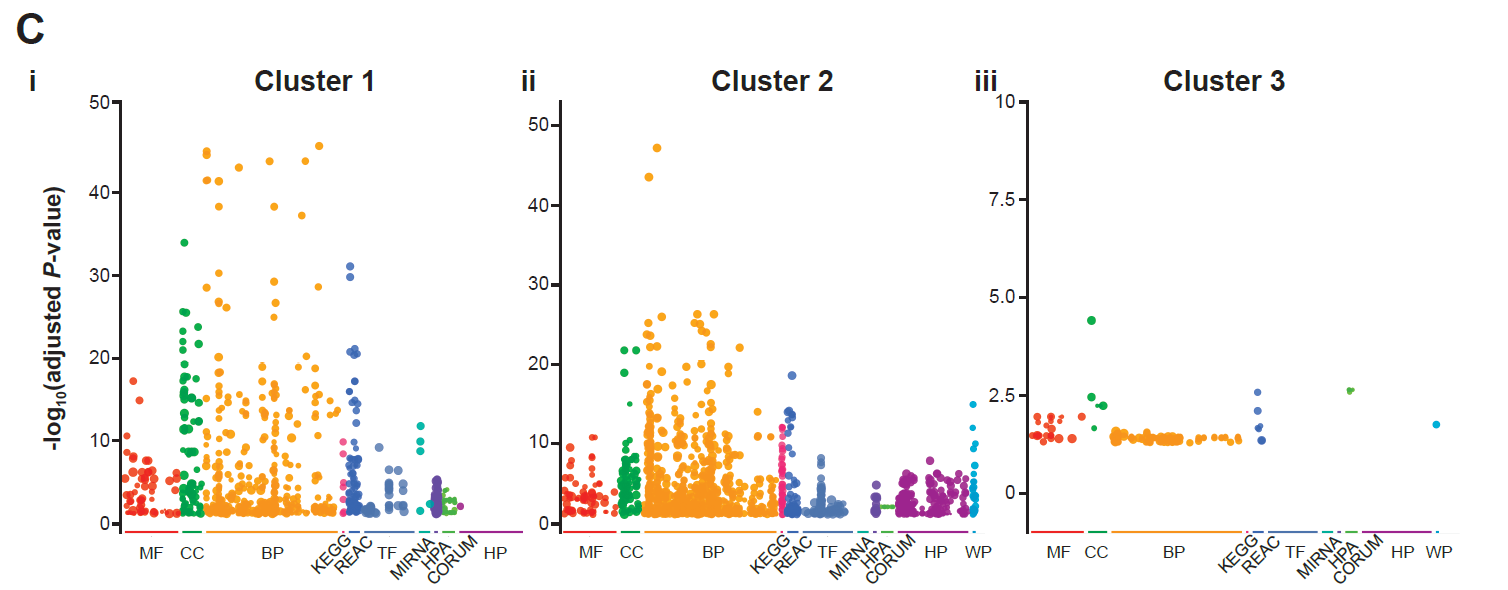

**D**

i

| Source | Term_name | Term_size | Query_size | Intersection_size | *P* value |
| --- | --- | --- | --- | --- | --- |
| REAC | Amplification of signal from the kinetochores | 92 | 50 | 14 | < 0.0001 |
| REAC | Amplification of signal from unattached kinetochores via a MAD2 inhibitory signal | 92 | 50 | 14 | < 0.0001 |
| REAC | Kinesins | 60 | 50 | 9 | < 0.0001 |
| REAC | Polo-like kinase-mediated events | 15 | 50 | 6 | < 0.0001 |

ii

| Source | Term_name | Term_size | Query_size | Intersection_size | *P* value |
| --- | --- | --- | --- | --- | --- |
| REAC | Interferon gamma signaling | 87 | 54 | 13 | < 0.0001 |
| REAC | Antigen Presentation: Folding, assembly and peptide loading of class I MHC | 25 | 54 | 9 | < 0.0001 |
| REAC | ER-Phagosome pathway | 90 | 54 | 12 | < 0.0001 |
| REAC | Endosomal/Vacuolar pathway | 11 | 54 | 7 | < 0.0001 |

iii

| Source | Term_name | Term_size | Query_size | Intersection_size | *P* value |
| --- | --- | --- | --- | --- | --- |
| REAC | Interleukin-10 signaling | 45 | 4 | 2 | 0.0026 |
| REAC | RUNX1 regulates transcription of genes involved in BCR signaling | 6 | 4 | 1 | 0.0186 |
| REAC | Interleukin-23 signaling | 9 | 4 | 1 | 0.0209 |
| REAC | NOTCH2 intracellular domain regulates transcription | 12 | 4 | 1 | 0.0223 |
| REAC | Signaling by NOTCH2 | 33 | 4 | 1 | 0.0436 |
